# Mindfulness meditators show altered distributions of early and late neural activity markers of attention in a response inhibition task

**DOI:** 10.1101/396259

**Authors:** NW Bailey, G Freedman, K Raj, CM Sullivan, NC Rogasch, SW Chung, KE Hoy, R Chambers, C Hassed, NT Van Dam, PB Fitzgerald

**Affiliations:** Monash Alfred Psychiatry Research Centre, Monash University Central Clinical School, Commercial Rd, Melbourne, Victoria, Australia; Brain and Mental Health Research Hub, School of Psychological Sciences, Monash Institute of Cognitive and Clinical Neurosciences, and Monash Biomedical Imaging, Monash University, Clayton, 3168 VIC, Australia; Campus Community Division, Monash University, Melbourne, Victoria, Australia; Department of General Practice, Monash University, Melbourne, Victoria, Australia; School of Psychological Sciences, The University of Melbourne, Parkville, VIC, Australia; Department of Psychiatry, Icahn School of Medicine at Mount Sinai, New York, NY, USA; Epworth Healthcare, The Epworth Clinic, Camberwell, Victoria, Australia, 3004

**Keywords:** Mindfulness Meditation, Electroencephalography, EEG, P3, C1, Go/Nogo, Response Inhibition, Attention

## Abstract

Attention is a vital executive function, since other executive functions are largely dependent on it. Mindfulness meditation has been shown to enhance attention. However, the components of attention altered by meditation and the related neural activities are underexplored. In particular, the contributions of inhibitory processes and sustained attention are not well understood. Additionally, it is not clear whether improvements in attention are related to increases in the strength of typically activated brain areas, or the recruitment of additional or alternative brain areas. To address these points, 34 meditators were compared to 28 age and gender matched controls during electroencephalography (EEG) recordings of neural activity during a Go/Nogo response inhibition task. This task generates a P3 event related potential, which is related to response inhibition processes in Nogo trials, and attention processes across both trial types. Compared with controls, meditators were more accurate at responding to Go and Nogo trials. Meditators showed a more frontally distributed P3 to both Go and Nogo trials, suggesting more frontal involvement in sustained attention rather than activity specific to response inhibition. Unexpectedly, meditators also showed increased positivity over the right parietal cortex prior to visual information reaching the occipital cortex. Both results were positively related to increased accuracy across both groups. The results suggest that meditators have an increased capacity to modulate a range of neural processes in order to meet task requirements, including higher order processes, and sensory anticipation processes. This increased capacity may underlie the improved attentional function observed in mindfulness meditators.

## Introduction

Attention is vital in selecting and maintaining processes most relevant for optimal behaviour (1). Attentional mechanisms have limited capacity and thus are most effective when allocated to processes that ensure behaviour consistent with the goals of the organism. In particular, attentional resources are most likely to enable optimal goal-oriented responses when the neural processes most at risk of failure are enhanced. In other words, attention improves goal-oriented behaviour by strengthening the weak links in the chain of neural processing that goes from stimulus detection to behavioural response (2, 3).

One method that enhances attention - mindfulness meditation - is conceptualised as a practice of training attention (or awareness) with an attitude of openness and non-judgement towards experiences (4, 5). Enhanced attention is a key mechanism of action in the improvements associated with mindfulness meditation (1, 5-9). Notably, meditators demonstrate improvements in sustained attention (10-12), distribution of scarce attentional resources in time (13, 14) and space (15), and attentional control including inhibition of prepotent behaviour (11, 16, 17). However, although eight-week standardized mindfulness programs improve aspects of cognition such as working memory and cognitive flexibility, they may not improve neuropsychological measures of attention (18). As such, individual components of attentional processes need further examination to determine the exact parameters of attentional function improvements result from mindfulness meditation.

Reviews suggest that mindfulness meditation most likely has its impact on attentional functions via changes to the structure and function of numerous regions in the prefrontal cortex, the anterior cingulate cortex, the insular cortex, and the hippocampus and amygdala (19, 20). As suggested above, sustained attention and inhibition are among the key mechanistic features from both an empirical and theoretical perspective (20). One task designed to test both inhibition and sustained attention is the Go/Nogo task. The Go/Nogo task presents stimuli to which participants are instructed to respond (Go trials), setting up a prepotent response tendency, and stimuli to which participants are instructed to withhold their response (Nogo trials). This task engages conflict monitoring to allocate neural resources between the two competing processes (response and non-response), keeping track of the alignment between behaviour (or potential behaviour) and the goals held by participants (21). Nogo trials also engage response inhibition to actively prevent a habitual or prepotent response (22). The Go/Nogo task also requires successful sustained attention, in order to keep track of stimuli, potential conflicts, and engage response inhibition processes (23). Improved behavioural performance on the Go/Nogo task has been shown after mindfulness meditation practice and was sustained for up to five months, reliably predicting improved socioemotional function (24).

At a neural level, sustained attention and inhibition are reflected by variations in the amplitude and synchronisation of neural oscillations, the average effect of which can be measured using event related potentials (ERPs) (25). Two ERPs are elicited by the Go/Nogo task: the N2, which is related to conflict monitoring and response inhibition, and significantly larger during Nogo trials (26-29) and the P3, which is thought to reflect attentional resource allocation, related to sustained attention on all trials (30, 31). Six studies have used the Go/Nogo task to examine the effect of trait mindfulness or mindfulness meditation on ERPs related to conflict monitoring, response inhibition, and sustained attention (see Table S1 for a summary). Each has studied a different population or intervention, and results between studies are inconsistent (32-37).

The inconsistencies make it difficult to draw meaningful conclusions about the effect of mindfulness meditation on attention. As the potential of meditation is likely to be most noticeable in those individuals who have engaged in extensive practice, work with this population is crucial to identifying likely benefits of mindfulness meditation. No such research to-date has examined neural response to the Go/Nogo task in long term meditators. Prior studies all used single electrode measures, further limiting potential conclusions. If meditation alters the P3 distribution, increasing prefrontal engagement (related to attention enhancements), single electrode analyses cannot differentiate these distribution differences from amplitude differences. The inconsistencies in prior studies may also be related to differences in windows and electrodes selected for analysis, and may have missed early processing changes that have been found in meditators in other tasks (15). An analysis technique encompassing all time windows and electrodes without a priori assumptions may be beneficial, in order to obtain a better understanding of the effect of meditation on neural activity. In particular, previous research has indicated that both voluntary and involuntary attention affects “evoked” sensory processing ERPs such as the C1, P1, N1, and P2 (38-41). Differences between meditators and controls in these windows are not detectable with research that focuses on typical Go/Nogo a priori windows of interest.

Recently developed EEG analytic techniques (42) enable comparison of neural activity across entire EEG epochs while simultaneously controlling family-wise error. Additionally, this analysis technique enables discrimination of differences reflecting altered overall neural response strength from changes in the distribution of neural activity across regions. This could elucidate whether meditation enhances the amplitude of typical neural responses related to sustained attention or inhibitory processes or trains a completely different pattern of brain region engagement, a question that has not been examined before in studies of meditation.

## Aims and Hypotheses

The aim of the current study was to assess whether individuals with extensive experience in mindfulness meditation showed differences in neural activity related to inhibition and sustained attention compared to demographically-matched individuals without meditation experience. We had hypotheses regarding both the amplitude and distribution of neural activity. Regarding amplitude, we hypothesized that: 1) neural activity related to conflict monitoring and response inhibition (the Nogo N2 and Nogo P3) would be larger in meditators, reflecting increased engagement of these neural processes as a result of the attention enhancing effect of meditation practice, and 2) neural activity related to attention would be larger in meditators (both Go and Nogo P3) reflecting increased engagement of these neural processes. Previous EEG research has not examined the distribution of neural activity independently of the amplitude of neural activity in meditators. However, research has suggested better attention and inhibition function are related to frontal activity (43, 44). As such, we hypothesised that the meditators would show more frontal activity in these ERPs, reflecting increased ability to engage the prefrontal cortex to maintain attention and inhibition processes. We also planned: 1) exploratory source analyses to assess which brain areas were activated during any topographical differences between groups and 2) microstate analysis to further characterise topographical differences between groups. Although simplified emotional faces were used as stimuli to replicate previous research using this task, no interaction between group and emotion was expected, as our previous research suggested the simplified faces were not sufficiently emotionally evocative to generate between group differences (45).

## Methods

### Participants and Self-Report Data

Thirty-six controls and 34 meditators were recruited through community advertising. Inclusion criteria for meditators involved a current meditation practice, with at least six months of meditation for at least two hours per week. All meditators except one had more than two years of meditation experience. Phone screening and in-person interviews were administered by experienced mindfulness researchers (GF, KR, NWB) to ensure meditation practices were mindfulness-based, using Kabat-Zinn’s definition - “paying attention in a particular way: on purpose, in the present moment, and nonjudgmentally” (46). Further screening ensured meditation practices were consistent with either focused attention on the breath or body-scan. Any screening uncertainties were resolved by between two researchers including the principal researcher (NWB). Control group participants did not have experience with meditation of any kind.

Exclusion criteria involved self-report of current or historical mental or neurological illness, or current psychoactive medication or recreational drug use. Participants were additionally interviewed with the MINI International Neuropsychiatric Interview for DSM-IV (47) and excluded if they met criteria for any DSM-IV psychiatric illness. Participants who scored in the mild above range or above in the Beck Anxiety Inventory (BAI) (48) or Beck Depression Inventory II (BDI-II) (49) were also excluded. All participants had normal or corrected to normal vision and were between 19 and 62 years of age.

Prior to completing the EEG task, participants reported their age, gender, years of education, handedness, and an estimate of the number of years spent meditating and the number of minutes per week spent meditating. Participants also completed the Freiburg Mindfulness Inventory (FMI) (50), Five Facet Mindfulness Questionnaire (FFMQ) (51), BAI, and BDI-II (see Table 1). All participants provided informed consent prior to participation. The study was approved by the Ethics Committee of the Alfred Hospital and Monash University (approval number 194/14).

**Table 1.** Demographic, self-report, and behavioural data.

|  | Meditators | Controls | Statistics |
| --- | --- | --- | --- |
|  | *M (SD)* | *M (SD)* |  |
| Age | 36.56 (10.88) | 35.68 (14.69) | $t(60) = 0.271, p = 0.794$ |
| Gender (F/M) | 21/13 | 17/11 | n.s. |
| Years of Education | 16.97 (2.55) | 15.87 (2.82) | $t(60) = 1.598, p = 0.115$ |
| Meditation Experience<br>(years) | 8.30 (10.28) | 0 |  |
| Current Time Meditating<br>Per Week (hours) | 5.50 (4.15) | 0 |  |
| BAI score | 4.24 (4.68) | 4.50 (5.62) | $t(60) = 0.202, p = 0.840$ |
| BDI score | 1.06 (1.87) | 1.61 (2.69) | $t(60) = 0.944, p = 0.349$ |
| FMI score | 45.62 (7.02) | 41.12 (7.75) | $t(60) = 2.401, p = 0.019^*$ |
| FFMQ score | 152.97 (17.13) | 138.39 (12.63) | $t(60) = 3.741, p < 0.001^{**}$ |

\* $p < 0.05$ , \*\* $p < 0.001$ .

Select data was excluded from analysis - four controls were excluded due to scoring in the mild depression range on the BDI, two due to misunderstanding task instructions, and one due to non-task completion. One additional control was excluded from neural analysis due to equipment fault. Two additional controls and three meditators were excluded from the behavioural analysis, due to an intermittent button fault during those sessions (enough correct response epochs were left for neural analysis, but accuracy calculations were insufficiently reliable). This left 28 controls for neural analysis (17 female, all right handed) and 27 controls for behavioural analysis. No exclusions were made for the meditators’ neural data, leaving 34 meditators (21 female, 3 left handed), and 31 for behavioural analysis.

### Task and Stimuli

Participants performed a Go/Nogo task with simplified emotional faces as stimuli while 64-channel EEG was recorded (see Figure 1). Task details were the same as Bailey et al. (45), with two blocks (instead of the four in the original design). The two blocks each included 75 happy and 75 sad faces. The equal trial type frequency was selected to limit between group comparisons to processes related to response inhibition (rather than also including processes related to probability of trial type, as would be the case if Nogo trials were less frequent than Go trials, since factors such as novelty modulate the Nogo N2 amplitude (52). Participants were instructed to respond by using both index fingers to press separate buttons simultaneously when they saw one emotion, and withhold responses to the other emotion. Stimulus-response pairings were reversed in the second block - participants who responded to happy faces in the first block responded to sad faces in the second block, and vice versa. Button press responses by the dominant hand were recorded.

**Figure 1.**
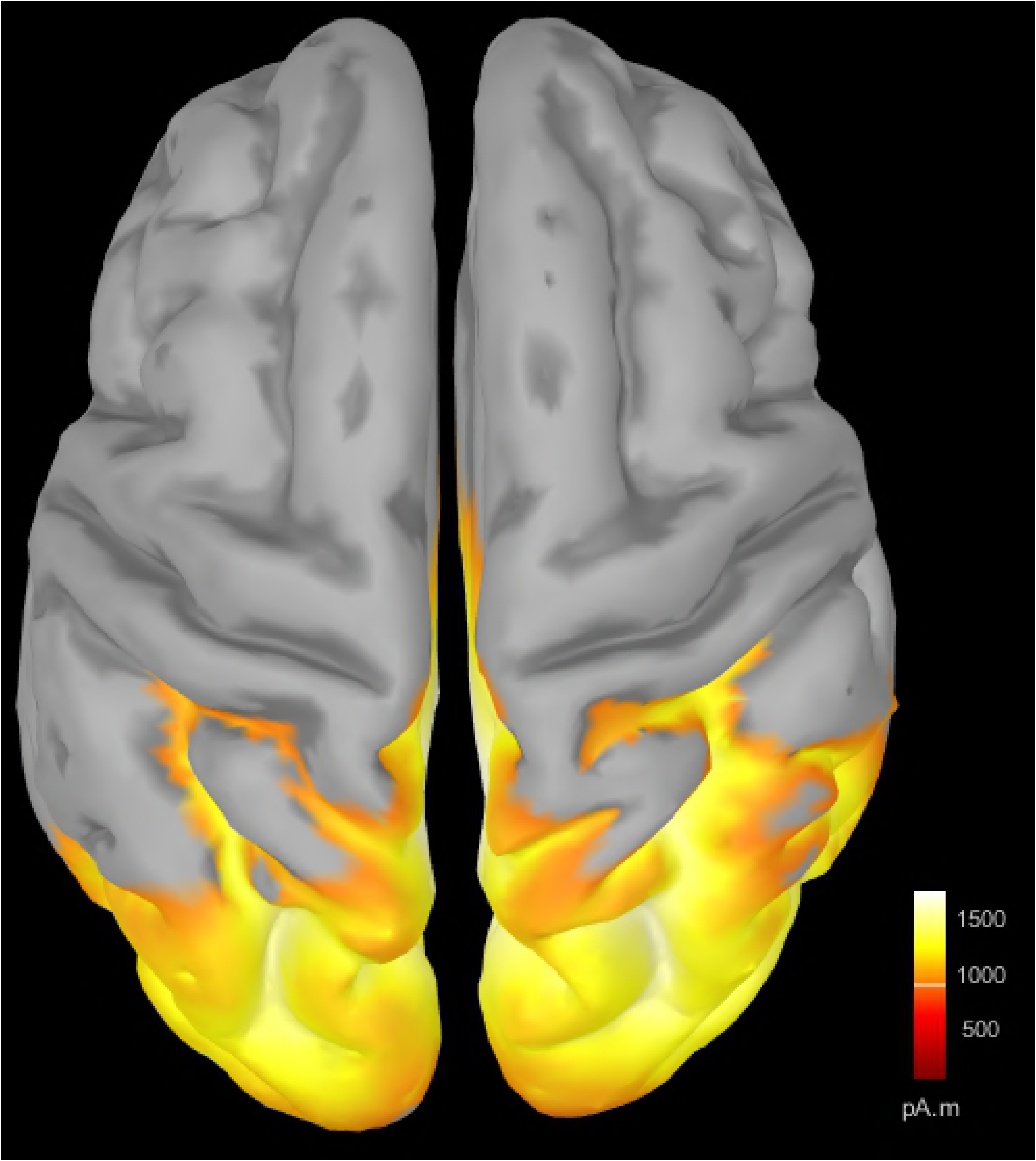
Go/Nogo task design. Go:Nogo ratio was 50:50, with stimulus response pairings switched in the second block so all participants responded to an equal number of happy and sad faces, and stimulus response pairings counter-balanced within each group.

Stimulus-instruction pairing was counterbalanced across participants and groups. Stimuli were presented for 250 ms, with an inter-trial interval of 900 ms (with a random jitter of 50 ms to avoid entrainment of EEG activity). Stimuli presentation was pseudo-random so that no more than four of each trial type was presented consecutively. Prior to beginning the task and again before the second block, participants were presented with a short practice block. The second practice was included to prevent extra errors and switching effects on the N2 amplitude (52). Percentage accuracy and reaction time (RT) for each trial type were extracted offline.

### Electrophysiological Recording and Pre-Processing

A Neuroscan 64-channel Ag/AgCl Quick-Cap was used to acquire EEG through NeuroScan Acquire software and a SynAmps 2 amplifier (Compumedics, Melbourne, Australia). Electrodes were referenced to an electrode between Cz and CPz. Eye movements were recorded with vertical and horizontal EOG electrodes. Electrode impedances were kept below 5kΩ. The EEG was recorded at 1000Hz, with an online bandpass filter of 0.1 to 100Hz.

Data were analysed offline in MATLAB (The Mathworks, Natick, MA, 2016a) using EEGLAB for pre-processing (sccn.ucsd. edu/eeglab) (53). Second order Butterworth filtering was applied to the data with a bandpass from 1–80 Hz and also a band stop filter between 47–53 Hz. Correct response trials were re-coded, and data were epoched from −500 to 1500 ms surrounding the onset of the stimulus presentation for each trial; only correct responses were analysed. Epochs were visually inspected by an experimenter experienced with EEG analysis and blinded to the group of each participant, and periods containing muscle artefact or excessive noise were excluded, as were channels showing poor signal. Thirty-five or more accepted epochs were obtained from each participant for each condition, and no significant differences were detected between groups in the number of accepted epochs (p > 0.10).

Data were combined with epoched data from another cognitive task (results of which will be presented in a separate publication) for Independent Component Analysis (ICA). AMICA (54) was used to manually select and remove eye movements and remaining muscle activity artefacts. Once artefactual ICA components were rejected, raw data were re-filtered from 0.1-80 Hz, all previous channel and epoch rejections were applied, and rejected ICA components were applied to this 0.1-80 Hz filtered data to avoid rejecting low frequency brain activity around 1 Hz (prior to ICA rejection, data below 1 Hz was filtered out as it adversely impacts the ICA process). Rejected electrodes were re-constructed using spherical interpolation (55). Data were then visually inspected again by a separate researcher (who was also blind to the group of the data inspected at that time) to ensure the artefact rejection process was successful. Recordings were re-referenced offline to an averaged reference and baseline corrected to the −100 to −10 ms period, and epochs from each condition and participant were averaged for ERP analyses.

### Source Localisation Pre-processing

Estimation of cortical sources during topographical between-group differences was performed using Brainstorm (56) (http://neuroimage.usc.edu/brainstorm/). EEG data were co-registered with the template model (ICBM 152) because individual MRIs were not available. The forward model used the Symmetric Boundary Element Method implemented in OpenMEEG software (57). The inverse model used the computation of minimum norm estimation, with sLORETA to normalise activity based on the depth of sources (58), with dipole orientations unconstrained to the cortex to minimize the impact of using the MRI template (59). Differences in estimation were calculated using absolute subtraction. We source localised the well-known P100 occipital ERP (averaged across 50 to 150 ms) to the correct location to demonstrate our source analysis was reliable even in the absence of individual MRI templates (see supplementary figure 1) (60). Statistical comparisons of source localisations were not performed, as scalp comparisons already demonstrated significant differences, and without MRI co-registration source statistical comparisons can be unreliable (61).

### Statistical Comparisons

Self-report and behavioural comparisons were made using SPSS version 23. Independent samples t-tests were conducted to ensure groups were matched in age, years of education, BAI, and BDI, and to determine whether groups differed in FMI, FFMQ scores. Chi square tests were used for gender and handedness. Percentage correct was compared with a repeated measures ANOVA involving 2 group x 2 Go/Nogo conditions x 2 emotion conditions. RT was compared in Go trials only (as these were the only trials requiring responses) with a repeated measures ANOVA involving 2 group x 2 emotion conditions. Fewer than 2 outliers were Winsorised for each percent correct condition. No outliers were present for cumulative percentage correct, and data met assumptions of normality and equality of variances. The Benjamini and Hochberg false discovery rate (FDR) (62) was used to control for multiple comparisons across behavioural performance measures.

### Primary Comparisons

Primary statistical comparisons for EEG data were conducted using the Randomised Graphical User Interface (RAGU) to compare scalp field differences across all electrodes and time points with randomisation statistics without making any a priori assumptions about electrodes or windows for analysis (42). This reference-free method takes advantage of the additive nature of scalp fields to allow comparisons of neural activity between groups and conditions without estimation of active sources by calculating a difference scalp field between groups or conditions. This difference scalp field shows the scalp field of brain sources that differed between the two groups/conditions, while brain sources that did not differ result in zero scalp field difference (42). RAGU controls for multiple comparisons in both time and space using randomisation statistics (see (42)). To control for multiple comparisons in time (which are made at each time point in the epoch), global duration statistics calculate the duration of significant effects that are longer than 95% of the significant periods in the randomised data, ensuring significant durations in the real data last longer than the random comparison data at p = 0.05 (42). Additionally, area under the curve statistics of significant time points across the entire epoch confirm sufficient control for multiple comparisons in the time dimension.

RAGU also allows for independent comparisons of overall neural response strength (with the global field power - GFP test) and distribution of neural activity (with the Topographic Analysis of Variance - TANOVA). Prior to the TANOVA, a Topographical Consistency Test (TCT) was conducted to ensure a consistent distribution of scalp activity within each group / condition. Lastly, Topographical Analysis of Covariance (TANCOVA) performs the same operations as TANOVA except it compares neural data to a linear predictor instead of between-group comparisons (42).

GFP and TANOVA tests were used to conduct 2 group x 2 Go/Nogo condition condition comparisons for averaged ERP data from −100 to 700 ms surrounding the onset of the stimulus. Five thousand randomisations were conducted with an alpha of p = 0.05. Post-hoc GFP and TANOVA tests to explore interactions were only conducted averaged across time periods of significant interaction after global duration controls.

In order to obtain effect sizes, GFP values were extracted from RAGU and submitted to parametric repeated measures ANOVA in SPSS. It is not feasible to compute traditional parametric effect sizes for topographical differences, the analysis of which includes many electrodes, so a randomisation statistic Cohen’s d equivalent was calculated for significant TANOVA differences. This method is reported in the supplementary materials.

The Benjamini and Hochberg false discovery rate (FDR) (62) was used to control for multiple comparisons for all comparisons testing primary hypotheses separately from comparisons involving behavioural data. FDR corrections were performed on the area under the curve p-values from each main effect or interaction. Area under the curve p-values were measured as the sum of all time points across the epoch in each comparison (across group main effects and group by Go/Nogo condition interaction). Controlling for multiple comparisons across both GFP and TANOVA tests, as well as across main effects and interactions avoided the hidden multiplicity in ANOVA designs (63). Post-hoc t-test designs were similarly controlled for using the FDR method. To enable comparison with other research, both corrected and uncorrected p-values are reported for significant comparisons (labelled ‘FDR p’ and ‘p-uncorrected’ respectively).

### Exploratory Analysis

Exploratory analyses were not corrected for multiple comparisons, so should be taken as preliminary findings. In order to assess relationships between behavioural results and neural activity, significant periods from group TANOVA comparisons were averaged and compared using TANCOVA tests with linear predictor values from significant between-group differences at the behavioural level.

Microstates are temporarily stable topographies of neural activation lasting approximately 80-120 ms before very quickly (∼5 ms) transitioning to another temporarily stable topography, reflecting difference source activations (64). Identification of microstates, determination of the optimal number of microstates, and statistical analysis was conducted using RAGU (65). Microstates were identified using atomize and agglomerate hierarchical clustering (AAHC) algorithm, which merges ERP topographics into clusters so that the average topography of the clusters explains maximal variance in the ERP (66). The optimal number of microstates was computed using cross-validation with the mean ERP from a learning set containing varied numbers of microstate classes and associated timing, which are then applied to the test set comprised of the remaining data. The optimal number of microstates is the point where the mean variance explained in the test set reaches its maximum (65). Randomisation statistics are then used to compare microstate properties during periods that were significant in the ERP TANOVA and GFP comparisons.

## Results

### Demographic and Behavioural

The neural analysis was the main focus of the study, so we only examined demographic and self-report differences for the participants included in the neural analysis. Results are summarised in table 1. For participants included in the neural analysis, no significant differences were present between groups in age, years of education, BAI score, BDI score, gender or handedness (all p > 0.3). Meditators showed significantly higher FMI t(60) = 2.401, p = 0.019 and FFMQ scores t(60) = 3.741, p < 0.001.

To examine behavioural performance, we compared percentage correct and reaction times. Normality, Box’s test, and Levene’s test were violated for percentage correct, however no significant interaction involving group was present with repeated measures ANOVA (Go/Nogo x group F(1,56) = 0.004, p = 0.952. Log10, natural log, and z-score transforms were attempted, but data remained non-normal. As such, corrections to normalise data were not performed. Cumulative percent correct across all conditions was calculated and found to be normally distributed. Meditators showed higher cumulative percentage correct with independent samples t-test t(56) = 2.511, p-uncorrected = 0.015 partial eta squared = 0.101, FDR p = 0.045.

No significant difference was found for any condition, group or interaction in the number of accepted epochs (all p > 0.10). No significant differences were found in reaction time for group comparisons or interactions involving group (all p > 0.10, see Table 2).

**Table 2.**
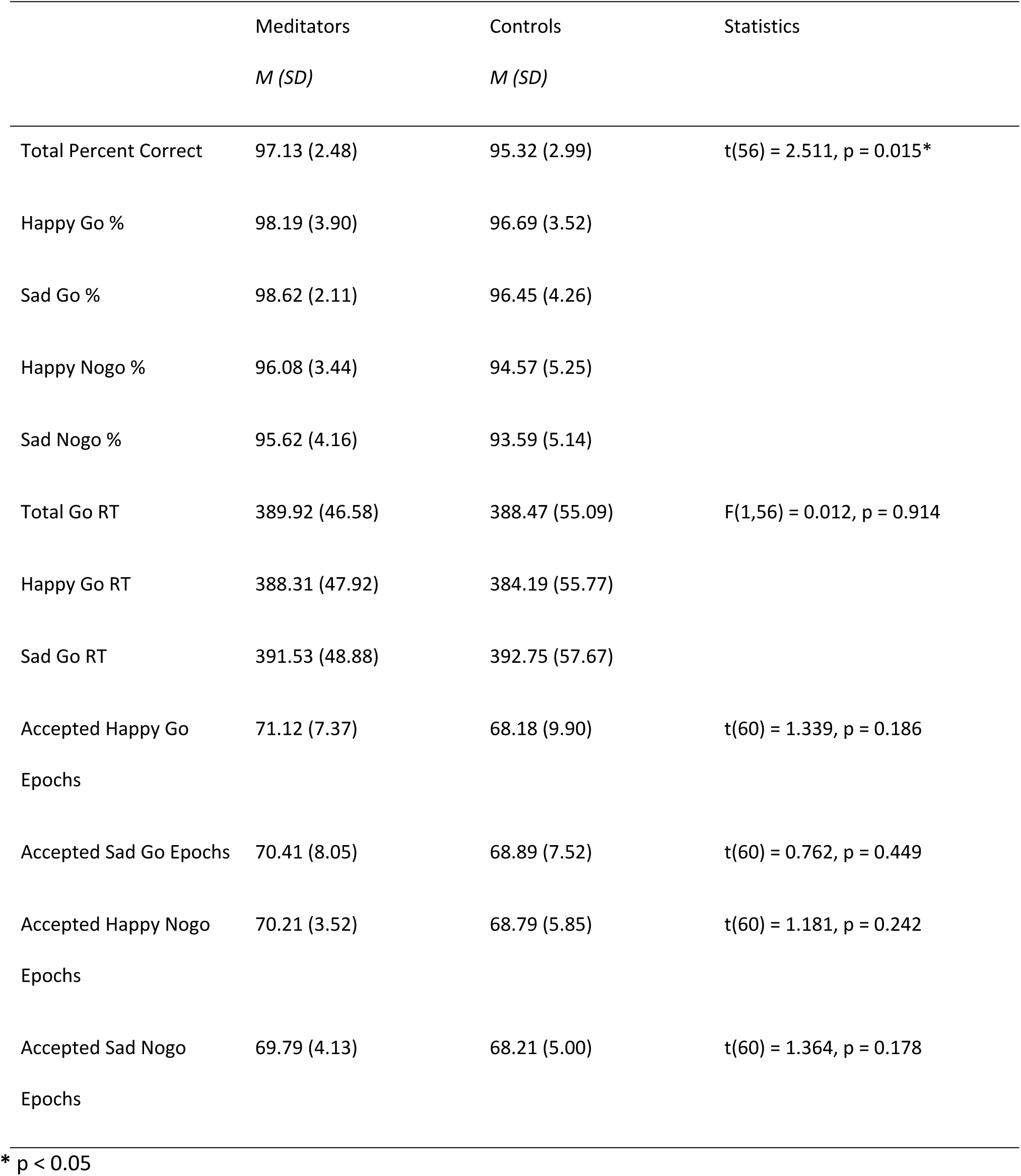
Behavioural and accepted epoch data.

Within the meditation group, no correlations were significant between cumulative percentage correct and meditation experience (years), minutes per week, or FFMQ scores (all p > 0.10).

## Neural Data

### Global Field Potential Test

To assess the strength of neural response to Go/Nogo trials, we analysed the data using the GFP test. A significant group by Go/Nogo trial interaction was present between 336 ms to 449 ms (area under the curve statistic p-uncorrected = 0.0198, FDR p = 0.0396), global duration statistic = 33 ms. When activity was averaged across the significant window (336 to 449 ms) to obtain a single value for analysis, the effect was still significant (p = 0.001). Post-hoc comparisons within trial type in RAGU indicated that controls and meditators did not differ in Go trial comparisons (p = 0.298) nor Nogo trial comparisons (p = 0.184). Controls showed a significant difference between Go and Nogo trials - Go trials showed larger amplitude than Nogo trials (p-uncorrected < 0.001, FDR p = 0.004). Meditators did not show a difference between Go and Nogo trial amplitudes (p = 0.743). See figure 2 for details. These results suggest that controls generate larger P3 amplitudes during Go trials, and smaller P3 amplitudes during Nogo trials, while meditators showed no differences. No differences were present in the N2 window (thought to reflect inhibition and conflict monitoring).

**Figure 2.**
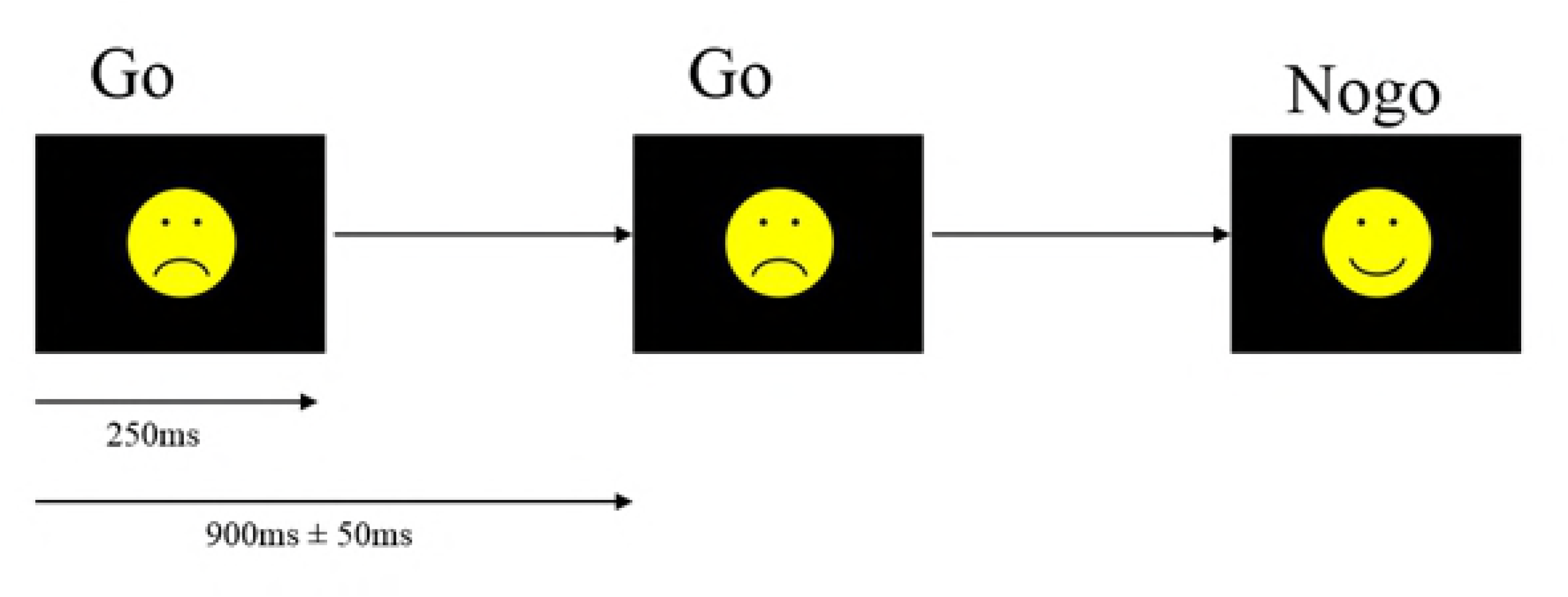
Significant group by Go/Nogo GFP interaction during the P3 window. A - Averaged GFP within the significant 336 ms to 449 ms window (green periods = 46 ms reflect periods that exceed the duration control for multiple comparisons across time = 33 ms). * p-uncorrected < 0.001 (FDR p < 0.004). B - Averaged topography during the significant window for each group. C – p-values of the group by Go/Nogo trial comparison for the real data against 5000 randomly shuffled permutations across the entire epoch.

In order to obtain effect sizes, GFP values were extracted from RAGU and submitted to a parametric repeated measures ANOVA in SPSS. Partial eta squared from Group x Go/Nogo interaction in parametric repeated measures ANOVA = 0.098. 95% Confidence intervals for controls Go = 2.253 to 2.999, Nogo = 1.718 to 2.349, meditators Go = 2.022 to 2.699, Nogo = 2.037 to 2.610. There was no main effect of group (p > 0.1).

### Topographical Consistency Test

In order to assess consistency of neural activity within groups and trial types, the TCT test was conducted (42). The TCT showed significant signal indicating consistency of neural activity within all groups / conditions across the entire epoch except prior to the stimulus and during a brief period (< 20 ms) at 550 ms in Nogo trials for controls, see figure 3). Consistent neural activity within conditions and groups indicates that TANOVA comparisons between conditions and groups are valid.

**Figure 3.**
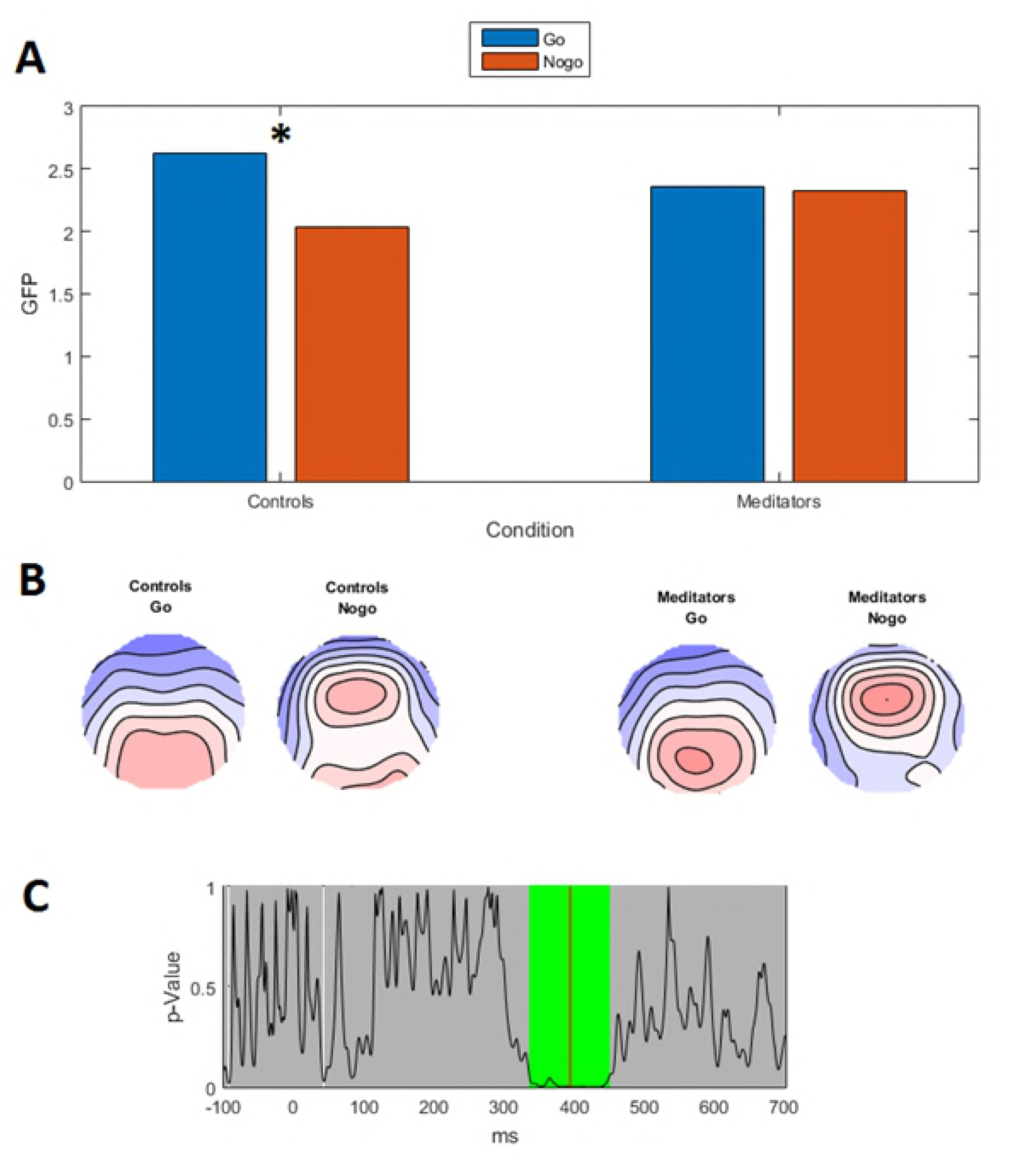
Topographical consistency test. The line indicates GFP values and the grey bars indicate p-values, with the red line indicating p = 0.05. White sections indicate regions without significantly consistent distribution of activity within the group/condition, while green periods indicate consistent distribution of activity across the group/condition after duration control for multiple comparisons across time (42). Note significant consistency across all conditions for both groups except for prior to stimulus onset, and around 550 ms in the Nogo trials for control participants.

### TANOVA

In order to examine potential differences in the distribution of neural activity in response to the Go and Nogo trials, TANOVAs were conducted. Significant main effects of group that survived duration control for multiple comparisons were present from −1 ms to 62 ms (prior to the C1 period, referred to as pre-C1 henceforth) (p = 0.003 averaged across the significant window, Cohen’s d equivalent effect size = 0.949), and from 416 ms to 512 ms (during the P3 period) (p = 0.007 averaged across the significant window, Cohen’s d equivalent effect size = 0.863). The area under the curve statistic for the entire epoch within the group main effect was p-uncorrected = 0.011 (FDR p = 0.040), and the global duration control statistic was 46 ms. Figures 4 and 5 depict the topographical differences between groups for the pre-C1 (−1 to 62 ms) and P3 (416 to 512 ms) periods respectively. No significant interaction between group and trial type was present (p > 0.1).

**Figure 4.**
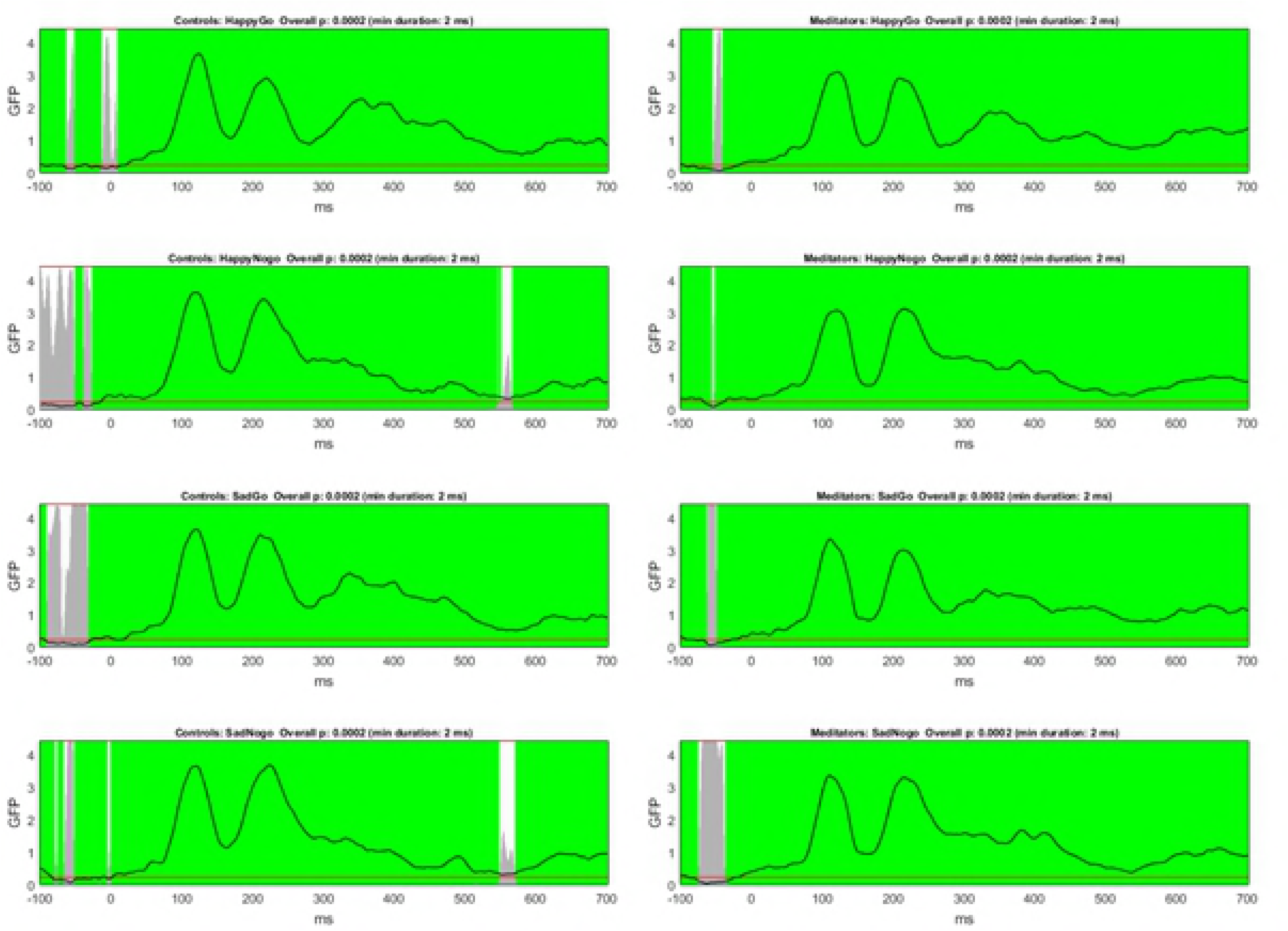
TANOVA main group effect from −1 to 62 ms. A - p values of the between-group comparison for the real data against 5000 randomly shuffled permutations across the entire epoch (green periods reflect periods that exceed the duration control for multiple comparisons across time = 46 ms). B - Averaged topographical maps for each group during the significant time window. C - p-map for meditators topography minus control topography during the significant time window (p = 0.003 averaged across the significant window, effect size = 0.9485).

**Figure 5.**
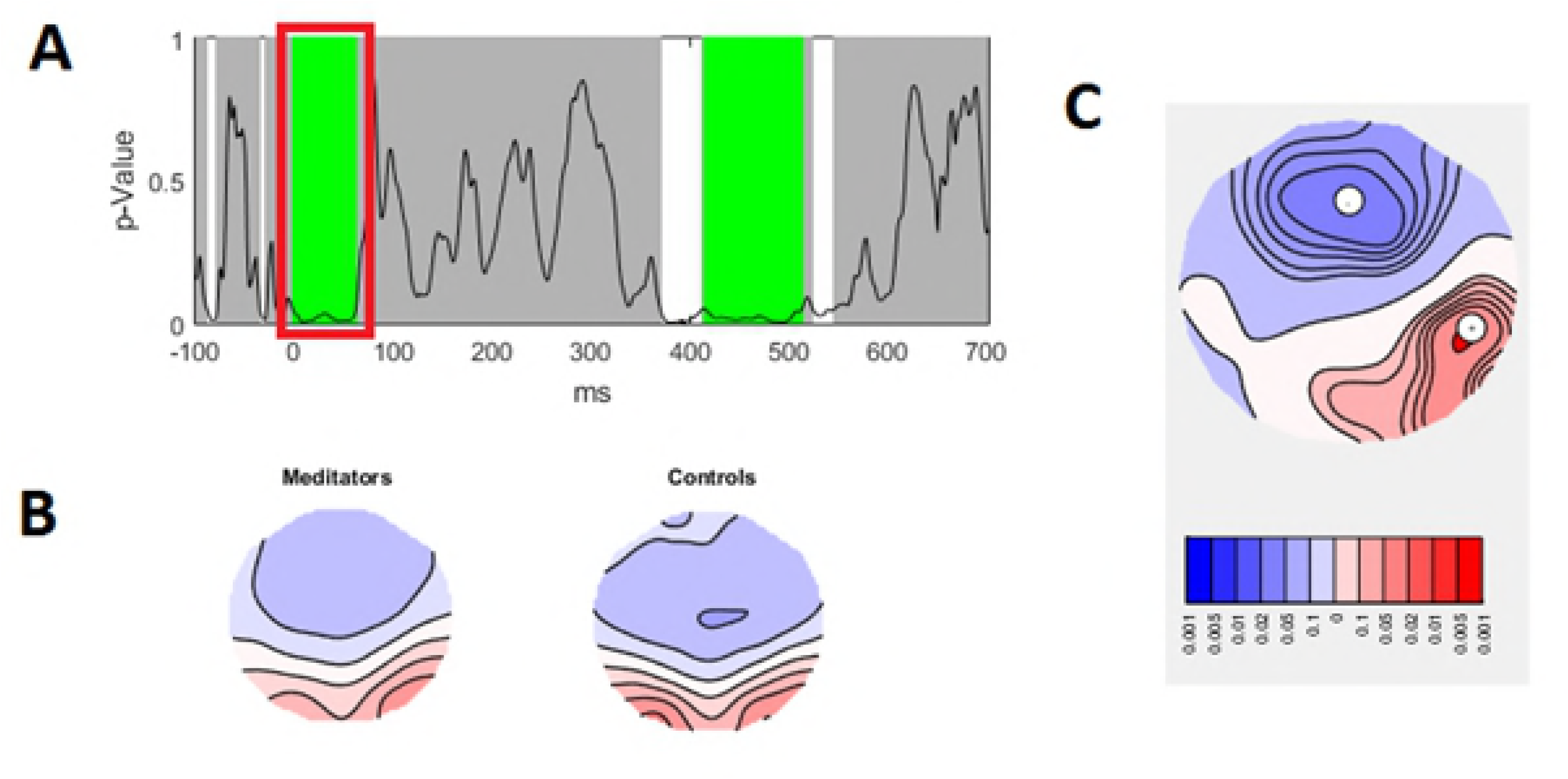
TANOVA main group effect from 416 to 512 ms. A - p values of the between-group comparison for the real data against 5000 randomly shuffled permutations across the entire epoch (green periods reflect periods that exceed the duration control for multiple comparisons across time = 46 ms). B - Averaged topographical maps for each group during the significant time window. C - p-map for meditators topography minus control topography during the significant time window (p = 0.007 averaged across the significant window, effect size = 0.863).

Overall, the differences indicate more fronto-central negativity and right posterior positivity in the meditation group during the pre-C1 (from −1 and 62 ms). Because the C1 is thought to be the first neural processing of visual stimuli (40, 67), the difference in pre-C1 activity is likely to reflect group differences in anticipatory activity.

The results also reflect more fronto-central positivity during the P3 in the meditation group. Because this difference was present across both Go and Nogo trials, the higher frontal activity in the meditation group may reflect altered attentional function of the P3 rather than altered inhibitory processes. No differences were present in the N2 window (thought to reflect inhibition and conflict monitoring).

### TANCOVA

To assess relationships between the altered distribution of neural activity shown by the TANOVA and behavioural performance, TANCOVAs were conducted between significant periods of activity in the TANOVA and cumulative percentage correct. Since participants performed at ceiling, groups were combined to maximise statistical power. TANCOVA between cumulative percentage correct and topographies averaged across the pre-C1 window (−1 to 62 ms) (p = 0.006) and the P3 window (416 to 512 ms) (p = 0.048) were significant, with positive topographics showing activity more similar to meditators, suggesting those topographies were related to better performance (see figure 6). However, this may be confounded by group differences in both topographies and performance. When running this analysis just within the meditation group the same pattern was apparent, but non-significant (p = 0.240 for −1 to 62 ms, and p = 0.766 for 416 to 512 ms), and the same was true for analysis within the control group (p = 0.112 for −1 to 62 ms, and p = 0.182 for 416 to 512 ms).

**Figure 6.**
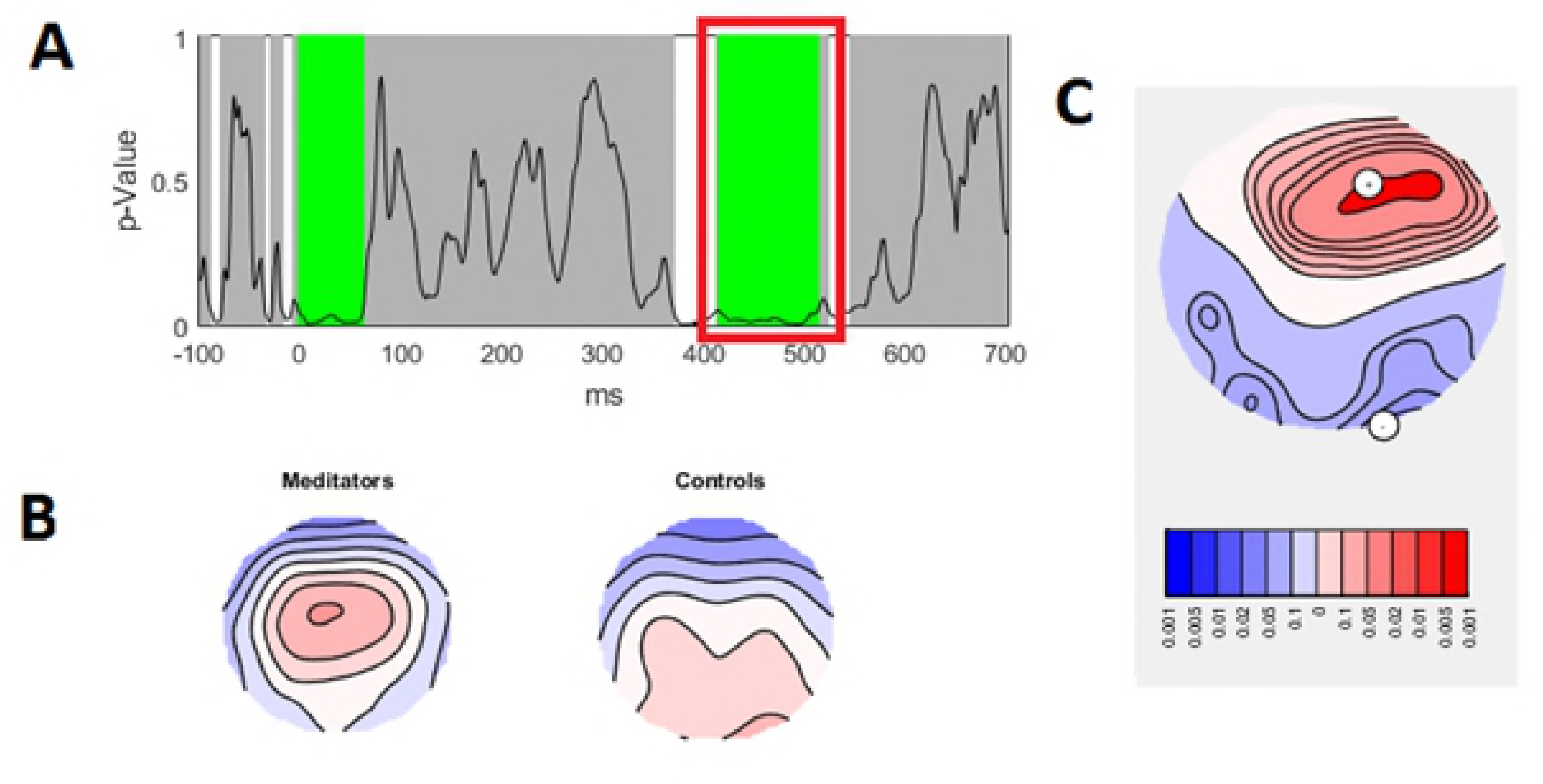
TANCOVA topographies depicting the relationship between cumulative percentage correct and averaged topography from −1 to 62 ms (left) and 416 to 512 ms (right) across both groups. * p = 0.048, ** p = 0.006.

### Microstates

To further explore the differences in ERPs, we used a microstate analysis approach which clusters different time periods into dominant scalp topographies. Microstate analyses were restricted to durations showing significant group main effects in the TANOVA (65). Three microstates differed in meditators – microstate 2, reflecting pre-C1 activity, and microstates 5 and 6, reflecting the P3 (see figure 7 for details). The findings suggested that pre-C1 period neural responses began earlier in meditators compared to controls, and that meditators spend more of the P3 period showing frontally dominant topographies compared to controls. Microstate 2 shows an earlier centre of gravity in meditators (the timepoint reflecting the centre of the GFP area for microstate 2 is earlier in meditators, p = 0.018), suggesting earlier processing of the stimuli in this group (the microstate is present from ∼0 to ∼100 ms following stimuli, matching TANOVA results in the −1 to 62 ms window). Microstate 5 shows a shorter duration in meditators (p = 0.003, meditators 78 ms, controls 217 ms). It also shows a smaller area under the curve in meditators (p = 0.031, meditators 105.9 ms x microvolts, controls 252.4 ms x microvolts), and an earlier centre of gravity (p = 0.028, meditators 318.2 ms, controls 381.9 ms). Microstate 6 shows more area under the curve in meditators (p = 0.044, 21.1 ms x microvolts in meditators, 0 ms x microvolts in controls). Microstate 5 is replaced by microstate 6 in meditators (indicating a more frontally distributed P3 during this period) but microstate 5 does not change to microstate 6 at all in controls. These results match the 416-512 ms period of significance in the TANOVA.

**Figure 7.**
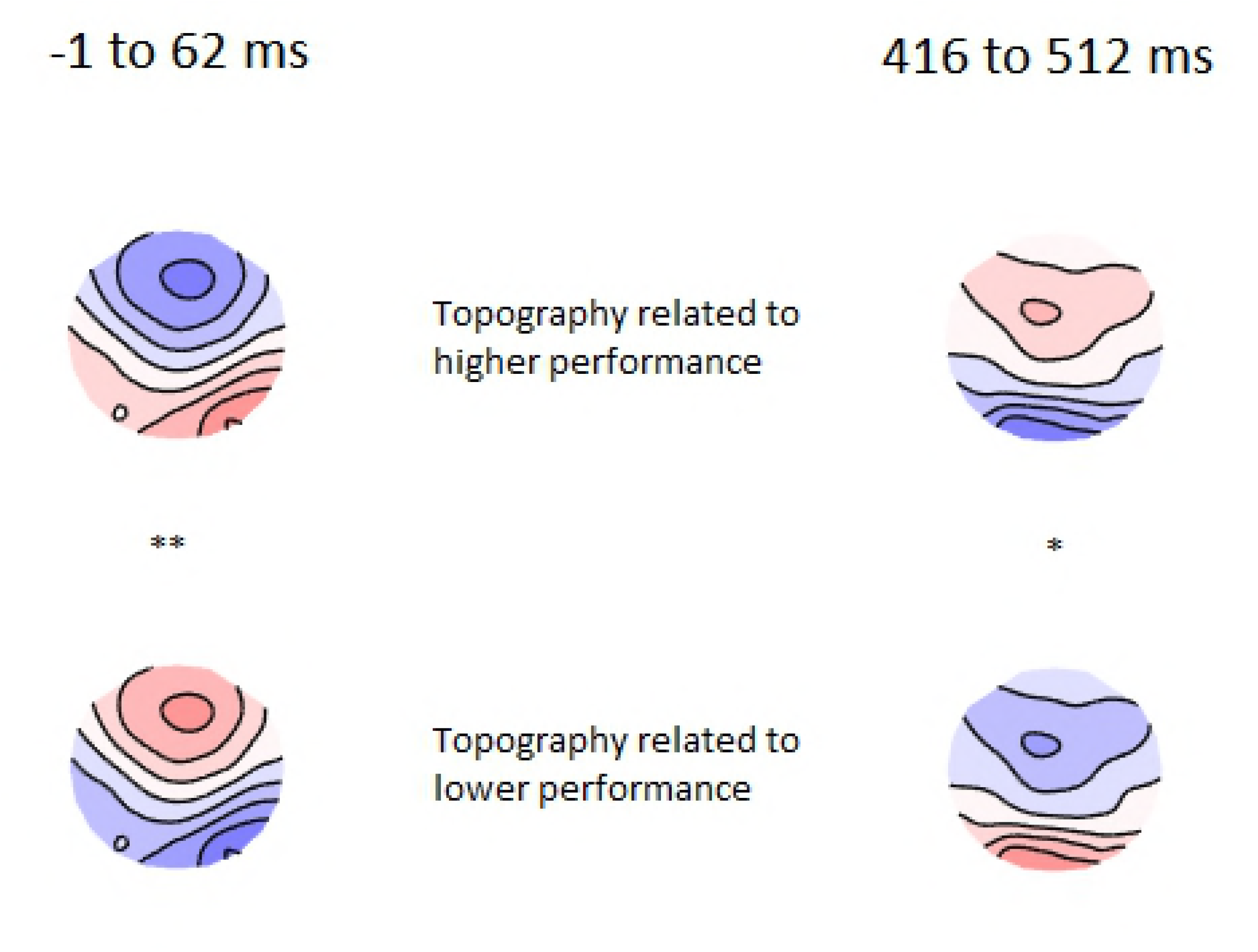
Microstate analysis showing overall between-group effects. Meditators differed in microstate 2 (reflecting pre-C1 activity), and microstates 5 and 6 (reflecting P3 activity). * p < 0.05 indicates an earlier centre of gravity in meditators, ** p < 0.01 indicates a longer duration in controls, + p < 0.05 indicates a larger area under the curve in controls, ^ p < 0.05 indicates larger area under the curve in meditators (6).

### Source analysis

To ascertain which brain regions contribute to the differences in scalp ERPs observed between the groups, we estimated the cortical sources of the signal using sLORETA. Source analysis suggested similar distributions of activity between the groups in both the pre-C1 and P3 time periods. Difference maps indicated that meditators showed more pre-C1 activity in right temporal and parietal regions, and a widespread pattern of more P3 activity in the central frontal and parietal regions. See figures 8 and 9 for details.

**Figure 8.**
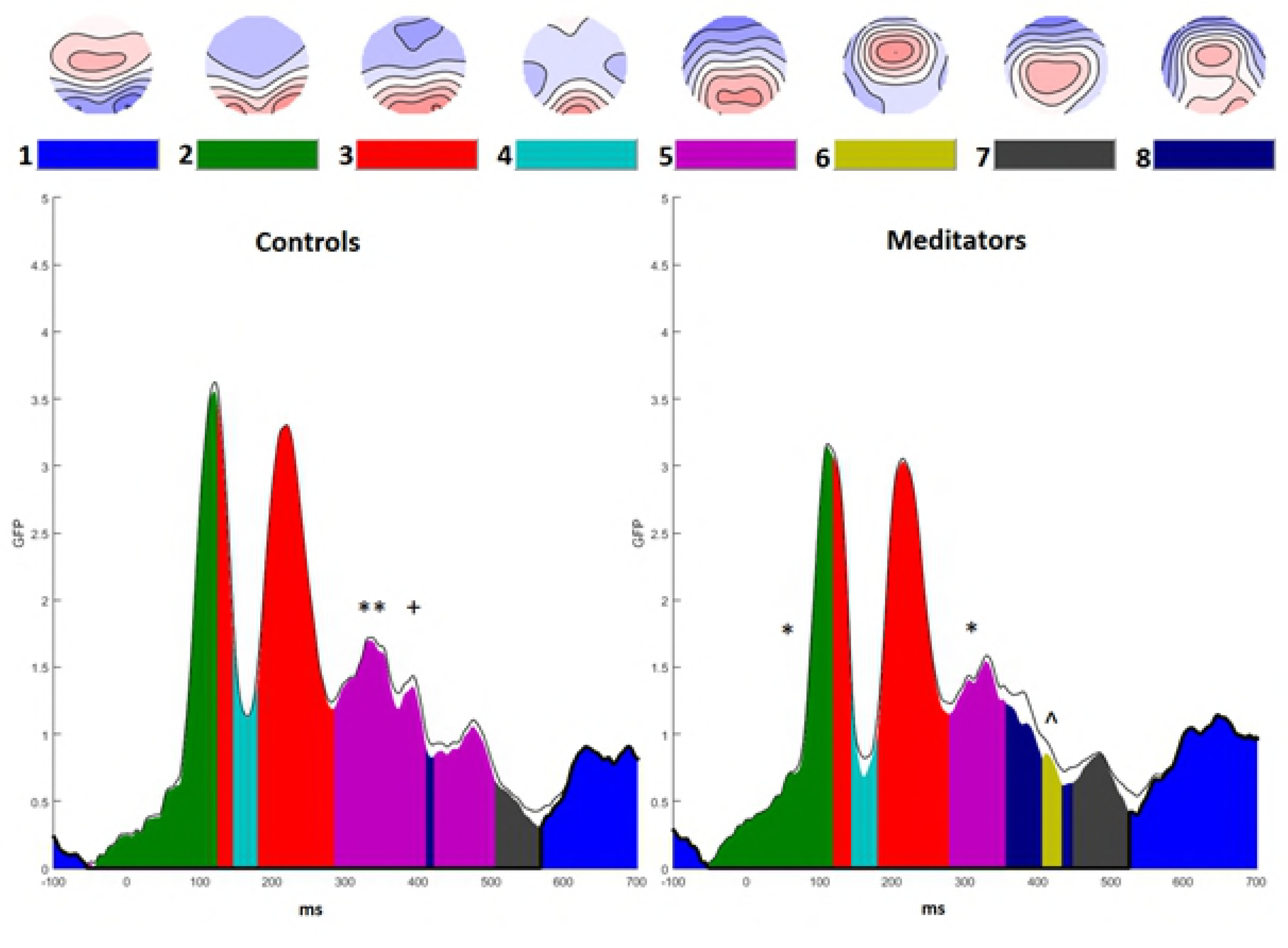
Source reconstruction during the −1 to 62 ms window using sLORETA and minimum norm imaging, unconstrained to cortex (to minimise assumptions). Group averages do not depict positive or negative voltages, only where a region was activated. Difference maps reflect meditator minus control activity (red reflecting more activity in meditators compared to controls, blue reflecting less activity in meditators).

**Figure 9.**
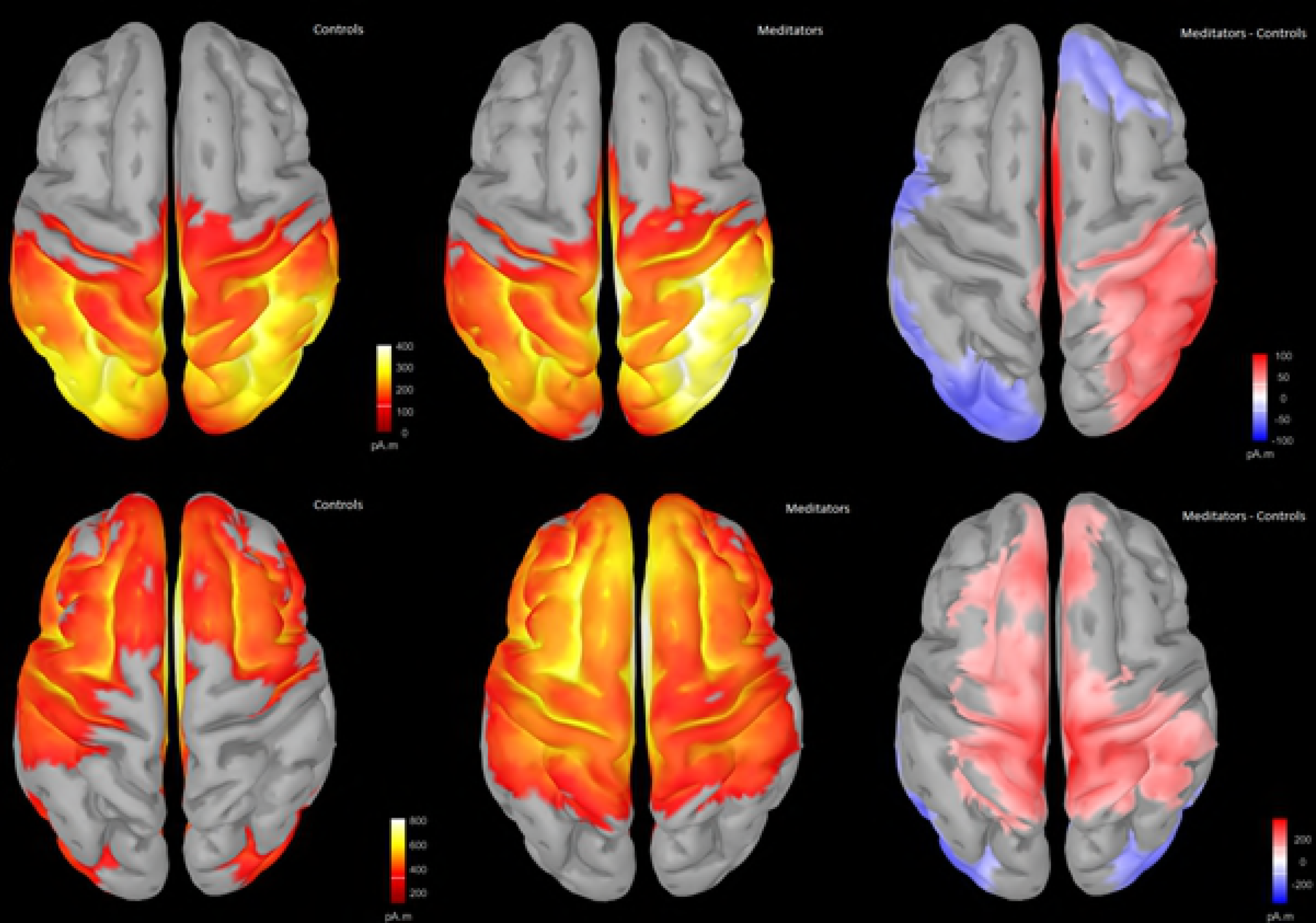
Source reconstruction during the 416 to 512 ms window using sLORETA and minimum norm imaging, unconstrained to cortex (to minimise assumptions). Group averages do not depict positive or negative voltages, only where a region was activated. Difference maps reflect meditator minus control activity (red reflecting more activity in meditators compared to controls, blue reflecting less activity in meditators).

## Discussion

Our study examined whether experienced mindfulness meditators showed differences in neural activity related to conflict monitoring, response inhibition, and sustained attention. The methods used enabled us to separately examine differences in the distribution of activated brain regions from differences in strength of neural activation, which has not been previously studied in meditators. Meditators showed higher accuracy across both Go and Nogo trials and frontally shifted distribution of neural activity during the P3 in both Go and Nogo trials. The latter finding suggests alterations to global attentional processes rather than inhibition specifically. Additionally, meditators showed less differentiation in the strength of neural activity between response and response inhibition trials during the early P3 window. Meditators showed more right parietal positivity during the pre-C1 period, suggesting more anticipatory neural activity for stimulus processing. The distribution of neural activity during both the pre-C1 and P3 significant time periods was correlated with behavioural performance across both groups, with better performing participants displaying the same distribution of activity observed in meditators.

These results suggest a range of differences in neural activity in mindfulness meditators compared to demographically matched controls. These differences likely reflect enhanced attentional mechanisms from long-term practice engaging attentional processes. The differences may reflect adaptive adjustments to the neural processes responsible for devoting resources to the functions maximally taxed by task requirements.

### A more frontally distributed P3

Meditators showed a more frontally distributed P3 than controls (Cohen’s d equivalent = 0.863). Previous research has indicated that engaging response inhibition processes generates a more frontally distributed P3, though no comparable work has explained the function of a more frontal P3 across both Go and Nogo trials. The more frontally distributed P3 in meditators across both trial types suggests that the differences were due to general attention effects rather than response inhibition. Additionally, source analysis indicated more activity in meditators across the superior/medial frontal gyrus, particularly the left hemisphere, as well as the bilateral parietal regions, spreading more laterally in the right hemisphere. Previous research has indicated the superior frontal gyrus to be related to executive function (44). Activity in the medial frontal gyrus is more pronounced when top-down control is allocated to Go/Nogo stimuli and is usually more related to attentional control than inhibition (43). Lastly, activity in the parietal cortex is thought to be related to selective and sustained attention, and the right parietal cortex to spatial attention (68, 69). These results suggest that the altered P3 distribution in meditators is likely to reflect enhanced attentional control. Further support for this conclusion comes from the finding that a more frontally distributed P3 was related to improved behavioural performance.

### Smaller P3 Amplitude Difference Between Response and Response Inhibition and No N2 Differences in Meditators

The meditation group showed no difference between Go and Nogo trials in overall neural response strength during the P3 window, while the control group showed larger neural response strength to Go trials than Nogo trials. However, no difference was found between groups in post-hoc comparisons of Go and Nogo trials independently, suggesting that trial type only differentiates neural response strength within controls rather than that meditators differ from controls. These results contradicted our expectation that the Nogo P3 would be enhanced in the meditation group, reflecting enhanced response inhibition (70). One potential explanation is that the task was easier for meditators. More difficult Go/Nogo tasks generate larger differences in neural activity between trial types (22). This explanation aligns with the better behavioural performance in the meditation group, who also showed less difference in neural activity between Go and Nogo trials.

Additionally, research has suggested that Go/Nogo ratios of 50/50 (as used in the current study) reduce response inhibition related neural activity by more than 60% (71). Equal ratio Go/Nogo tasks may simply compare general response-related activity to trials where response-related activity is never initiated (rather than trials that initiate response activity that subsequently must be inhibited (71)). As such, observed differences may merely reflect improved attentional control in meditators rather than improved inhibitory processes. In support of this explanation, studies using harder Go/Nogo tasks with lower frequencies of Nogo trials show enhanced Nogo P3 activity in ADHD patients who have undergone meditation training (33). However, the Go and Nogo P3 is influenced by stimuli frequency (72). Thus, future research examining response inhibition in meditators should attempt to differentiate between frequency and response inhibition effects.

Additionally, although an interaction between trial type and group was present for P3 amplitudes, no interaction was present in the behavioural data. The lack of behavioural difference likely reflects a ceiling effect – the behavioural results were too consistently high to reveal an interaction, while neural data may be more sensitive. Lastly, we expected the N2 component to be altered in meditators. Previous research with infrequent Nogo trials has demonstrated N2 alterations from meditation (32, 36). As mentioned with the lack of altered Nogo P3 activity in meditators, the N2 component may not have differentiated the groups because response inhibition processes were not sufficiently taxed by the equiprobable Go/Nogo task.

### More Right Posterior Pre-C1 Positivity

The meditators showed a topography with more negative fronto-central activity and more right posterior positivity during the pre-C1 window (Cohen’s d equivalent = 0.9485). The result reflects differences in neural activity that precedes the earliest point that visual related activity has been shown to reach the occipital cortex (∼ 50 ms post stimulus (40, 67)). Meditators showed differences in neural activity *before* stimulus perception. Anticipatory activity is present during periods leading up to stimulus processing, reflecting top-down attentional control to enhance cortical processing of stimuli, ensuring optimal processing (73-76). In other words, the pre-C1 may reflect enhanced endogenous attention, which has been defined as *“the exercising of an intention to selectively attend, based on some internal representation of what will be attentionally relevant in the near future. This intention interacts with attention deployment systems to reorganize the attentional set of the brain in preparation for incoming stimuli—a preparatory attentional state*” (77).

Such anticipatory activity has been found in the dorsal visual processing stream (in temporo-occipital regions), with top-down influences from lateral parietal attentional gating regions, and frontal control regions (67, 73, 77). These regions overlap with those shown in our source analysis. These areas may exert an excitatory effect on primary visual areas that increase and prolong stimulus processing in those areas (78). Thus, this pre-C1 difference may reflect an improved attentional preparedness among meditators, reflecting a greater readiness for stimulus processing and enhanced focus, as is a goal in many early mindfulness meditation practices (4, 6-8, 11).

Additionally, the right occipital and temporal regions have been shown to specialise in processing faces as well as for anticipation of general visual processing, suggesting that higher activity in these regions in the meditation group is likely to assist stimuli processing in the current task (77, 79). As such, the results could reflect enhancement of the visual processing pathway so the chain of information from perception to performance is more effective (67). Although unexpected, our pre-C1 results provide further evidence for the suggestion that enhanced attention in meditators reflects a stronger ability to modulate neural activity towards the optimal achievement of goals (2, 80). As such, this difference in anticipatory pre-C1 activity may reflect an altered top-down brain state that prepares meditators’ brains for the subsequent perceptual brain states. This may have enabled an increased ability of meditators to sustain attentional focus on the chosen object and by consequence to show enhanced behavioural performance (which in this case are the task stimulus) (7).

These results have clinical implications - research indicates that aversive stimuli cause altered visual processing within 60-120 ms (81). Individuals with anxiety also show stronger neural responses to negative emotional images within the 80 ms C1 period (82). This early response to aversive stimuli and early over-activation in anxious individuals reflects early sensory processing bias that may be impossible for the higher order functions to later modulate. The clinical benefit of mindfulness may involve alteration to attentional mechanisms that allow modulation of early neural processing, reducing emotional reactivity before emotional reactions are elicited. This may explain why mindfulness has amongst its strongest clinical effects on anxiety (83).

### Strengths, Limitations and Future Directions

Although a strength of the current study is the selection of a well-matched control group, the main limitation is the lack of ability to draw conclusions about causation. It may be that individual differences such as personality factors that predispose that group towards mindfulness meditation are ultimately responsible for the differences. Previous longitudinal research has indicated that mindfulness meditation does alter neural activity (13, 14, 17, 84) partially mitigating such concerns. Nonetheless, it is difficult to control for potential self-selection biases among those who have chosen to meditate versus those who have not (Davidson & Kaszniak, 2015). A parsimonious and robust interpretation of the current conclusions (and those of other cross-sectional studies of experienced meditators) is that differences relate to “leading a life that involves meditation” but the research offers no information as to whether meditation is causal in the differences.

Another strength of the current study is the confirmation that both the meditation and control groups showed consistent topographical activation patterns prior to performing between-group comparisons. This is important, because it eliminates the possibility that differences in within-group variability could explain between-group differences, despite absence of signal within one of the groups (because the signal was variable in that group and averaged out to zero).

Future research would do well to examine the commonalities and differences between altered neural activity in mindfulness meditators across different tasks. This is necessary to answer questions about whether the neural effects of mindfulness meditation are process-driven or domain-specific (9). Our suggestion is that the changes that result from meditation reflect enhancement no of one specific neural process, but of the modulation of a range of oscillatory activity, in order to strengthen the weakest link in the chain of neural processes. As such, we would expect that the process most pressured by a specific task may demonstrate enhanced function in meditators who have improved attentional function. We recommend including easy and hard conditions for research comparing meditators to controls. This would enable identification of neural processes that are upregulated to enable performance in the hard condition, allowing determination of whether that process is specifically affected by enhanced attention in mindfulness meditators.

### A Propositional Integrative Interpretation

Overall, the results show differences in both anticipation of sensory processing and top down attention related differences in neural activity in mindfulness meditators, in alignment with previous research (13, 14, 17, 84). The altered topographies suggest that different neural assemblies are recruited in meditators to perform the same task but with increased accuracy, rather than the same neural assemblies being more strongly activated.

We suggest that the differences in meditators reflect improved attentional function, and this improved attentional function provides enhancements to neural processes that are the ‘weakest link’ in achieving task-oriented goals (2, 80, 85). Attention supports the processes most likely to fail in the chain from stimulus processing to response, reducing the chance of failure at those most vulnerable points and enhancing the probability of successful task performance.

In this context, differences between meditators and controls are likely to be task-specific rather than neural activity or region-specific. Other tasks are likely to demonstrate different effects from meditation depending on the neural processes most taxed by the task, for example alpha modulation enhancements to reduce somatosensory distraction (86), or theta synchronisation to stimulus in attentional blink tasks (14). Indeed, the current sample of meditators showed an alternative profile of differences compared to controls than the differences found in the current study when they performed both a colour and emotional Stroop task (Raj et al. in preparation) and an N-back task with a tactile distractor (Freedman et al. in preparation). This interpretation may provide an explanation for the variation in findings between studies comparing meditators to controls, as different neural processes are engaged by different tasks and varied task parameters. We hope that the results of the current study can be interpreted and contextualised within this framework, in combination with future research, to provide a more sophisticated understanding of how neural activity differs in meditators.

## Acknowledgements

PBF has received equipment for research from MagVenture A/S, Medtronic Ltd, Cervel Neurotech and Brainsway Ltd and funding for research from Neuronetics and Cervel Neurotech. PBF is on the scientific advisory board for Bionomics Ltd. All other authors have no conflicts to report. PBF is supported by a National Health and Medical Research Council of Australia Practitioner Fellowship (6069070). KEH is supported by a National Health and Medical Research Council of Australia Career Development Fellowship (1082894). NCR is supported by a National Health and Medical Research Council of Australia Fellowship (1072057).

## Supplementary Materials

The following is a summary of previous research examining mindfulness with EEG activity during the Go/Nogo task. Six studies have used the Go/Nogo task to examine the effect of trait mindfulness or mindfulness meditation on ERPs related to conflict monitoring, response inhibition, and sustained attention (see Table 1 for a summary). Each has studied a different population or intervention, and results between studies are inconsistent (32-37). Studies have included healthy university students trained in one week of mindful deep breathing (32), adult ADHD and depression provided with MBCT for twelve weeks (33-35), healthy adolescents trained in eight weeks of the.b foundations mindfulness program (36), and healthy university students measuring the relationship between trait mindfulness and neural activity (37). All studies showed larger N2 amplitudes in the mindful participants, suggesting conflict monitoring or response inhibition processes. However, each study showed N2 alterations to different trial types, with (32) showing the increase in infrequent Nogo trials (and only in the 5 minutes / day condition), (36) to frequent Nogo trials, and (33) to frequent Go trials (but not Nogo trials), and (37) to both trial types (in individuals showing higher trait mindfulness). This suggests inconsistency in the relationship between neural changes resulting from mindfulness and conflict monitoring (low frequency trials) and response inhibition processes (Nogo trials). In addition to the inconsistency regarding N2 changes, only Schoenberg et al and Quaglia et al (33, 37) showed increased Nogo P3, suggesting enhanced response inhibition closure in more mindful participants. No study showed overall P3 changes (related to sustained attention), which we would expect from mindfulness meditation (which effects sustained attention).

**Table S1.**
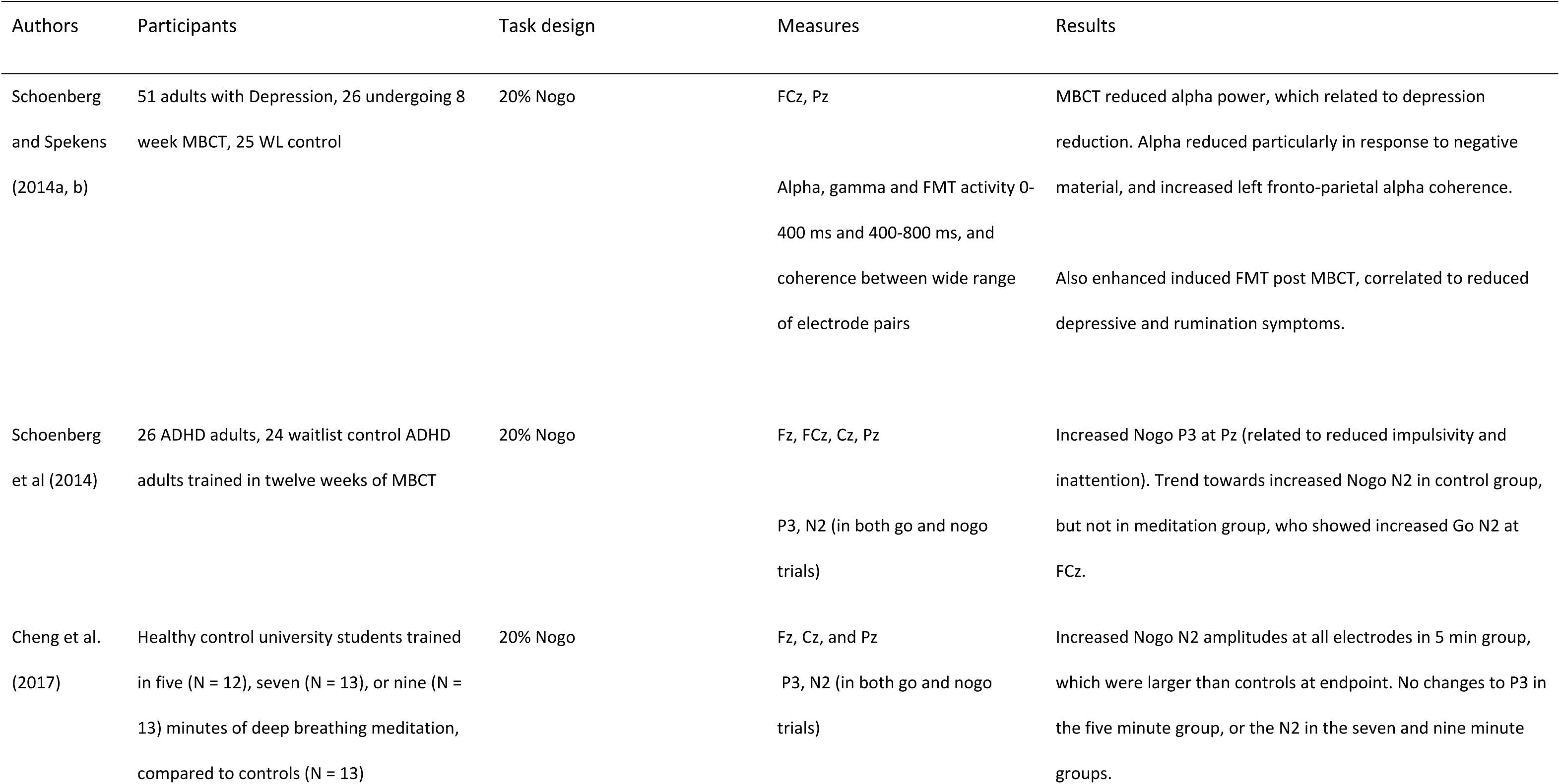
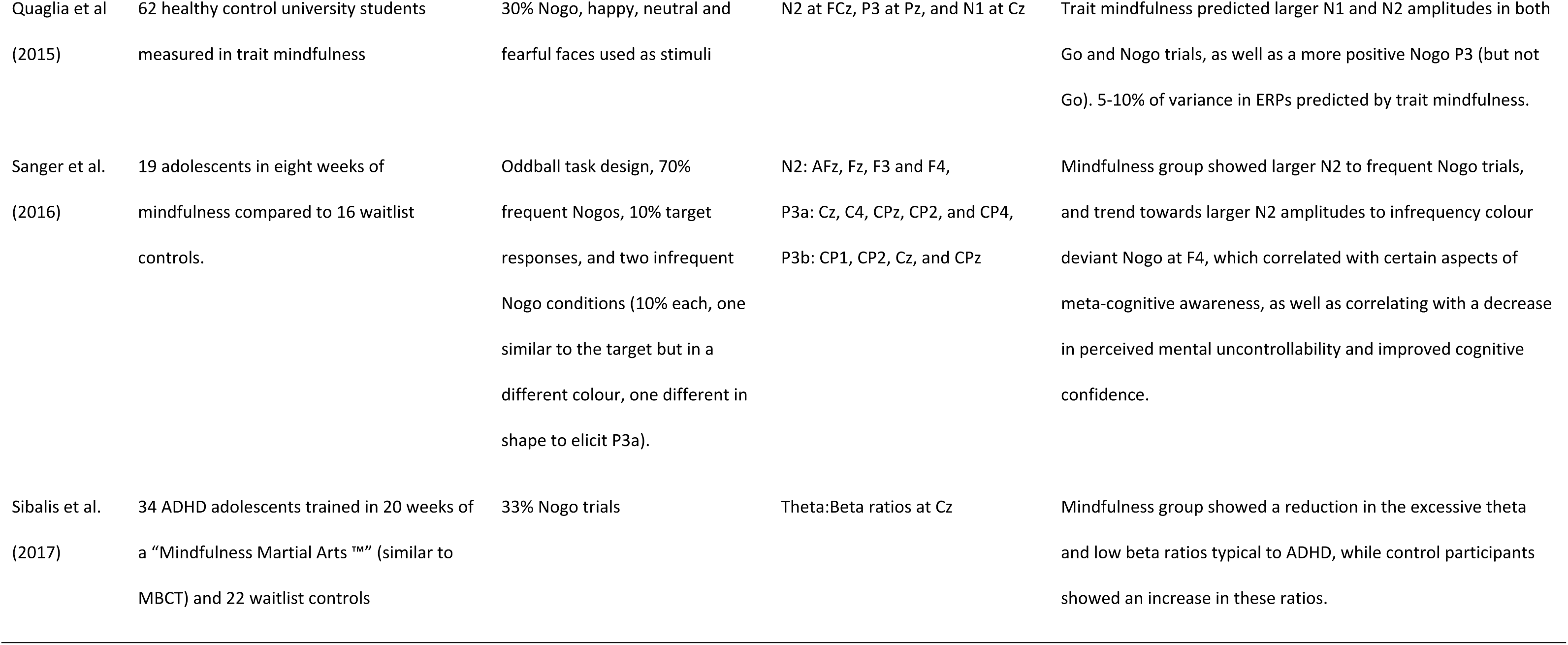
Previous mindfulness research using the Go/Nogo task

| Authors | Participants | Task design | Measures | Results |
| --- | --- | --- | --- | --- |
| Schoenberg and Spekens (2014a, b) | 51 adults with Depression, 26 undergoing 8 week MBCT, 25 WL control | 20% Nogo | FCz, Pz<br>Alpha, gamma and FMT activity 0-400 ms and 400-800 ms, and coherence between wide range of electrode pairs | MBCT reduced alpha power, which related to depression reduction. Alpha reduced particularly in response to negative material, and increased left fronto-parietal alpha coherence.<br>Also enhanced induced FMT post MBCT, correlated to reduced depressive and rumination symptoms. |
| Schoenberg et al (2014) | 26 ADHD adults, 24 waitlist control ADHD adults trained in twelve weeks of MBCT | 20% Nogo | Fz, FCz, Cz, Pz<br>P3, N2 (in both go and nogo trials) | Increased Nogo P3 at Pz (related to reduced impulsivity and inattention). Trend towards increased Nogo N2 in control group, but not in meditation group, who showed increased Go N2 at FCz. |
| Cheng et al. (2017) | Healthy control university students trained in five (N = 12), seven (N = 13), or nine (N = 13) minutes of deep breathing meditation, compared to controls (N = 13) | 20% Nogo | Fz, Cz, and Pz<br>P3, N2 (in both go and nogo trials) | Increased Nogo N2 amplitudes at all electrodes in 5 min group, which were larger than controls at endpoint. No changes to P3 in the five minute group, or the N2 in the seven and nine minute groups. |
| Quaglia et al.<br>(2015) | 62 healthy control university students<br>measured in trait mindfulness | 30% Nogo, happy, neutral and<br>fearful faces used as stimuli | N2 at FCz, P3 at Pz, and N1 at Cz | Trait mindfulness predicted larger N1 and N2 amplitudes in both<br>Go and Nogo trials, as well as a more positive Nogo P3 (but not<br>Go). 5-10% of variance in ERPs predicted by trait mindfulness. |
| Sanger et al.<br>(2016) | 19 adolescents in eight weeks of<br>mindfulness compared to 16 waitlist<br>controls. | Oddball task design, 70%<br>frequent Nogos, 10% target<br>responses, and two infrequent<br>Nogo conditions (10% each, one<br>similar to the target but in a<br>different colour, one different in<br>shape to elicit P3a). | N2: AFz, Fz, F3 and F4,<br>P3a: Cz, C4, CPz, CP2, and CP4,<br>P3b: CP1, CP2, Cz, and CPz | Mindfulness group showed larger N2 to frequent Nogo trials,<br>and trend towards larger N2 amplitudes to infrequency colour<br>deviant Nogo at F4, which correlated with certain aspects of<br>meta-cognitive awareness, as well as correlating with a decrease<br>in perceived mental uncontrollability and improved cognitive<br>confidence. |
| Sibalis et al.<br>(2017) | 34 ADHD adolescents trained in 20 weeks of<br>a “Mindfulness Martial Arts <sup>TM</sup>” (similar to<br>MBCT) and 22 waitlist controls | 33% Nogo trials | Theta:Beta ratios at Cz | Mindfulness group showed a reduction in the excessive theta<br>and low beta ratios typical to ADHD, while control participants<br>showed an increase in these ratios. |

## Supplementary Methods

### Statistical methods

RAGU’s GFP test uses the spatial standard deviation of the electric field to compare the global strength of cortical activation (42). The TANOVA assesses the percentage of randomly shuffled data sets that show larger scalp field differences between groups / conditions than the real data to determine whether to accept / reject the null hypothesis at a predetermined alpha level (42). Prior to the TANOVA, a Topographical Consistency Test (TCT) was conducted, comparing global field power within each group / condition to randomly shuffled data to ensure a consistent distribution of scalp activity within each group / condition. A significant TCT test confirms that any potential differences in the TANOVA are due to actual group / condition differences, rather than simply high variation within one of the groups (87). Because each of these tests only use a single value for comparison between groups / conditions at each time point (the spatial standard deviation for the GFP test, and the scalp field difference for the TANOVA and TCT test), no controls for multiple comparisons in the spatial dimension are needed even though all electrodes are included. To control for multiple comparisons in time (which are made at each time point in the epoch), global duration statistics calculate the duration of significant effects that are longer than 95% of the significant periods in the randomised data, ensuring significant durations in the real data last longer than the random comparison data at p = 0.05 (42). Additionally, global count statistics and area under the curve statistics of significant time points were checked to confirm sufficient control for multiple comparisons in the time dimension. The recommended L2 normalisation of scalp field variance across sensors was administered to remove scale differences, so that significant results in the TANOVA reflect a different distribution of neural activity without being affected by amplitude (42).

Because the measure that TANOVA compared to the random permutations in order to calculate statistical comparisons was the overall difference between groups in the summed standard deviation of the GFP from each electrode, traditional effect size calculations (which require measures from each individual) were not feasible. In consultation with the author of RAGU, TANOVA effect sizes were calculated using an analogue of Cohen’s d from z-scores of the topography differences between groups (Koenig, personal communication, 2017). Firstly, the real data difference between groups in topographical distribution of activity was extracted for each time point during the significant period, then averaged across the period (RDm). Secondly, the mean difference score from each time point across all permutations were calculated, and the mean (PDm) and SD (PDsd) of this across time permutation mean were calculated. The permutation mean was then subtracted from the real data mean, and this value was divided by the SD to obtain a z-score. This z-score was then divided by the square root of (the proportion of the total number of participants from the first group (P1) multiplied the proportion of the total number of participants from the second group (P2), multiplied by the total number of participants (N) minus 2). This final value is an analogue of Cohen’s d.

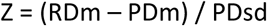

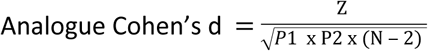

## Supplementary Results

Supplementary figure 1. Source reconstruction during the well-known P100 occipital ERP, averaged across the 50 to 150 ms window across both groups using sLORETA and minimum norm imaging, unconstrained to cortex (to minimise assumptions). This was performed to demonstrate our source analysis was reliable even in the absence of individual MRI templates (60). Note that the average does not depict positive or negative voltages, just whether a region was activated.

## Supplementary Discussion Points

Anticipatory activity has been found in the dorsal visual processing stream (in temporo-occipital regions), with top-down influences from lateral parietal attentional gating regions, and frontal control regions (67, 73, 77). These regions are similar to those shown in our source analysis. Additionally, attentional manipulations have been found from 0 to 50 ms time locked to stimuli presentation (85). Previous research examining these anticipatory brain responses and our current results cumulatively indicate that differences were not present in the primary visual cortex, suggesting attentional mechanisms modulating sensory processing, rather than altered initial sensory processing (77). These areas may exert an excitatory effect on primary visual areas that increase and prolong stimulus processing in those areas (78).

Additionally, the right occipital and temporal regions have been shown to specialise in processing faces as well as for anticipation of general visual processing, suggesting that higher activity in these regions in the meditation group is likely to assist stimuli processing in the current task (77, 79). As such, the results could reflect enhancement of the visual processing pathway so the chain of information from perception to performance is more effective (67). However, if the enhanced anticipation explanations for the altered pre-C1 topography are accurate, it is not clear why the difference detected in the pre-C1 period did not lead to alterations during the C1 and later sensory processing ERPs (despite being related to improved behavioural performance and enhanced theta synchronization to stimuli during the P3). We suspect neural activity for the time spanning the pre-C1 period to the P3 (including the C1, P1, N1, P2, and N2) must differ between meditators and controls, but in a manner too subtle to detect using ERP analyses (perhaps reflected by frequency or connectivity changes, or in action potentials too small to be detected at the scalp level). Future research may able to offer further insights.

The pre-C1 differences also raise a question. If the pre-C1 differences reflect altered anticipatory activity in meditators, do the results conflict with the conception of mindfulness as “being in the present moment”? Anticipatory activity necessitates processing ahead of the present moment. It may be that the concept of “being in the present moment” is a higher order neuropsychological processes, while automatic processes such as the C1 may still be engaged to anticipate stimuli, even as a result of the higher order processes resting “in the present moment”. In this explanation, non-judgemental awareness may apply not to discrete moments as measured with the millisecond precision of ERPs, but across the second or so that we feel subjectively aware, such that expectation for a stimuli within that second is “seeing reality as it is”. In which case, the conflict is between conceptual levels of processing, and essentially reflects a technical conflict. An alternative explanation is that the altered pre-C1 reflects lack of expectation, allowing neural processes during the pre-C1 window to be more available for processing the stimuli. Or one last alternative - activity in this region is simply strengthened in general in meditators, and as such the pre-C1 differences do not reflect anticipation or reaction to the stimuli, but just that the area is more prone to being active. However, the area was not activated more during other periods of the epoch, so the initial explanations seem more sensible to us.

An interesting and potentially useful feature of C1 activity is that, due to the anatomical structure of the primary visual cortex, the C1 polarity reverses depending on whether visual stimuli are presented in the upper or lower visual field (40). Future research could determine whether the altered pre-C1 in meditators reflects anticipation by presenting a visual processing task to meditators frequently in the lower visual field, and unexpectedly and infrequently in the upper visual field. If the effect of meditation on the pre-C1 reflects increased anticipation, unexpected stimuli may show a disruption of stimulus processing reflected by a reduced C1 compared to controls. In contrast, if the effect is a result of lack of expectation, the unexpected change in stimulus location will engage the C1 more quickly or strongly in meditators than controls (similar to the task tested by Kelly et al (40)).

## Comparisons with previous research

Our results were different to previous research using the Go/Nogo task. Previous research has shown increased amplitude of Nogo P3 (33) and increased Nogo N2 (32, 36), or decreased Nogo N2 (33). The current research showed less differentiation between P3 amplitudes to Go and Nogo trials in the meditation group, and a more frontal P3 topography. However, as mentioned, previous research used a lower proportion of Nogo trials, setting up an increased prepotent response tendency, likely exerting different task demands and different neural weak points for attentional improvements in the meditation group to focus upon. Additionally, while the current research found topographical differences during the pre-C1 and P3 period, previous research performed analyses using single electrode analyses, so could not discriminate between topography and amplitude changes, and during specified windows not including the pre-C1 period. Lastly, previous research examined less experienced meditators as participants, and included ADHD (33, 88), depressed (34, 35), and adolescent participants (36), so would have obtained smaller, perhaps less consistent and stable differences between groups, perhaps reflecting more short term changes to brain activity rather than permanent trait-like changes, and differences reflecting different preceding neural profiles of clinical and younger populations (for example reversal of the typical excess theta reduced beta ratio in ADHD (88)). Also, as mentioned in the introduction, the previous research is inconsistent.

More broadly, the current results align with a plethora of research indicating enhanced attention in mindfulness meditators. These studies have also shown that multiple aspects of attentional function are improved, including sustained attention (10-12), distribution of scarce attentional resources in time (13, 14) and space (15), and attentional control including inhibition of prepotent behaviour (11, 16, 17).

Lastly, as well as discriminating differences in strength of neural response from distribution, no previous research has examined the consistency of neural activity within meditation groups. Given the variability in practice, individuals, and other unknown factors, differences between meditation and control groups may in fact reflect simply differences in the degree of variability between groups. Traditional parametric statistics cannot account for within-group consistency (42). Demonstrating consistent within-group activity for meditators is an important step in ensuring the neural changes are practice-induced, and that the changes are common across the group. While not feasible to study longitudinally, prolonged meditation practice over many years is likely to result in the most consistent, durable and significant effects, and studying these individuals is likely to provide stronger conclusions about the effects of mindfulness meditation (9). As such, the consistent neural activity within groups in the current research is a strength of the study.

## Additional Potential Limitations

It is important to note that potential explanations for the function of the altered mechanisms in meditators are based on correlational research. As such, we cannot be certain that the mechanism performs the suggested function, simply that it is related to it. A demonstrative example is that although the strongest correlation with hippocampal theta in rats is walking behaviour, complete lesioning of the hippocampus does not alter walking behaviour (89).

Because the pre-C1 effect was unexpected and has not been shown by prior research, we are less confident that it reflects a real difference between groups. However, it did show a large effect size and was highly significantly related to performance. The result should be replicated and explored further by future research, in order to confirm and explain the finding in more detail.

Another limitation is that our results may not generalise to clinical populations in which mindfulness is most commonly used - the mechanism of action in treatment interventions may be different. While mindfulness meditation is likely to alter attentional mechanisms in clinical groups, it may be that the practice does not alter the same mechanisms in those individuals as it does in healthy controls. As per our integrated interpretation of our results, attentional training is likely to alter neural activity where alterations are most necessary to improve function. These may be different in clinical and healthy populations. For example, depressed participants show altered Go/Nogo N2 activity compared to controls (45), so it may show activity in the N2 window that is altered in depressed individuals who practice mindfulness meditation.

Lastly, it should also be noted that although the language used in this study may suggest an agency behind attentional function, attention is not a homunculus guiding neural activation to achieve the goals it has set. The psychological concept of attention *is* the action of the neural activity, and other neural processes self-organising to achieve “goals”, which are themselves also simply behavioural reflections of neural processes. The origins of consciousness have not yet been explained.

## References

1. Posner MI, Rothbart MK, Tang Y-Y. Enhancing attention through training. Current Opinion in Behavioral Sciences. 2015;4:1–5.

2. Lavie N. Perceptual load as a necessary condition for selective attention. Journal of Experimental Psychology: Human perception and performance. 1995;21(3):451.

3. Lubbe RHVd, Vogel RO, Postma A. Different effects of exogenous cues in a visual detection and discrimination task: delayed attention withdrawal and/or speeded motor inhibition? Journal of Cognitive Neuroscience. 2005;17(12):1829–40.

4. Crane RS, Brewer J, Feldman C, Kabat-Zinn J, Santorelli S, Williams JMG, et al. What defines mindfulness-based programs? The warp and the weft. Psychological medicine. 2017;47(6):990–9.

5. Malinowski P. Neural mechanisms of attentional control in mindfulness meditation. Frontiers in neuroscience. 2013;7:8.

6. Hölzel BK, Lazar SW, Gard T, Schuman-Olivier Z, Vago DR, Ott U. How does mindfulness meditation work? Proposing mechanisms of action from a conceptual and neural perspective. Perspectives on psychological science. 2011;6(6):537–59.

7. Lutz A, Slagter HA, Dunne JD, Davidson RJ. Attention regulation and monitoring in meditation. Trends in cognitive sciences. 2008;12(4):163–9.

8. Shapiro SL, Carlson LE, Astin JA, Freedman B. Mechanisms of mindfulness. Journal of clinical psychology. 2006;62(3):373–86.

9. Slagter HA, Davidson RJ, Lutz A. Mental training as a tool in the neuroscientific study of brain and cognitive plasticity. Frontiers in human neuroscience. 2011;5:17.

10. Chambers R, Lo BCY, Allen NB. The impact of intensive mindfulness training on attentional control, cognitive style, and affect. Cognitive Therapy and Research. 2008;32(3):303–22.

11. Moore A, Malinowski P. Meditation, mindfulness and cognitive flexibility. Consciousness and cognition. 2009;18(1):176–86.

12. Valentine ER, Sweet PL. Meditation and attention: A comparison of the effects of concentrative and mindfulness meditation on sustained attention. Mental Health, Religion & Culture. 1999;2(1):59–70.

13. Slagter HA, Lutz A, Greischar LL, Francis AD, Nieuwenhuis S, Davis JM, et al. Mental training affects distribution of limited brain resources. PLoS biology. 2007;5(6):e138.

14. Slagter HA, Lutz A, Greischar LL, Nieuwenhuis S, Davidson RJ. Theta phase synchrony and conscious target perception: impact of intensive mental training. Journal of cognitive neuroscience. 2009;21(8):1536–49.

15. Van Leeuwen S, Singer W, Melloni L. Meditation increases the depth of information processing and improves the allocation of attention in space. Frontiers in human neuroscience. 2012;6:133.

16. Manna A, Raffone A, Perrucci MG, Nardo D, Ferretti A, Tartaro A, et al. Neural correlates of focused attention and cognitive monitoring in meditation. Brain research bulletin. 2010;82(1-2):46–56.

17. Moore AW, Gruber T, Derose J, Malinowski P. Regular, brief mindfulness meditation practice improves electrophysiological markers of attentional control. Frontiers in human neuroscience. 2012;6:18.

18. Lao S-A, Kissane D, Meadows G. Cognitive effects of MBSR/MBCT: a systematic review of neuropsychological outcomes. Consciousness and cognition. 2016;45:109–23.

19. Fox KC, Nijeboer S, Dixon ML, Floman JL, Ellamil M, Rumak SP, et al. Is meditation associated with altered brain structure? A systematic review and meta-analysis of morphometric neuroimaging in meditation practitioners. Neuroscience & Biobehavioral Reviews. 2014;43:48–73.

20. Tang Y-Y, Hölzel BK, Posner MI. The neuroscience of mindfulness meditation. Nature Reviews Neuroscience. 2015;16(4):213.

21. Botvinick MM, Cohen JD, Carter CS. Conflict monitoring and anterior cingulate cortex: an update. Trends in cognitive sciences. 2004;8(12):539–46.

22. Benikos N, Johnstone SJ, Roodenrys SJ. Varying task difficulty in the Go/Nogo task: the effects of inhibitory control, arousal, and perceived effort on ERP components. International Journal of Psychophysiology. 2013;87(3):262–72.

23. Barkley RA. Behavioral inhibition, sustained attention, and executive functions: constructing a unifying theory of ADHD. Psychological bulletin. 1997;121(1):65.

24. Sahdra BK, MacLean KA, Ferrer E, Shaver PR, Rosenberg EL, Jacobs TL, et al. Enhanced response inhibition during intensive meditation training predicts improvements in self-reported adaptive socioemotional functioning. Emotion. 2011;11(2):299.

25. Klimesch W, Sauseng P, Hanslmayr S, Gruber W, Freunberger R. Event-related phase reorganization may explain evoked neural dynamics. Neuroscience & Biobehavioral Reviews. 2007;31(7):1003–16.

26. Donkers FC, Van Boxtel GJ. The N2 in go/no-go tasks reflects conflict monitoring not response inhibition. Brain and cognition. 2004;56(2):165–76.

27. Falkenstein M. Inhibition, conflict and the Nogo-N2. Clinical Neurophysiology. 2006;117(8):1638–40.

28. Falkenstein M, Hoormann J, Hohnsbein J. ERP components in Go/Nogo tasks and their relation to inhibition. Acta psychologica. 1999;101(2-3):267–91.

29. Nieuwenhuis S, Yeung N, Cohen JD. Stimulus modality, perceptual overlap, and the go/no‐go N2. Psychophysiology. 2004;41(1):157–60.

30. Datta A, Cusack R, Hawkins K, Heutink J, Rorden C, Robertson IH, et al. The P300 as a marker of waning attention and error propensity. Computational intelligence and neuroscience. 2007;2007.

31. Wickens C, Kramer A, Vanasse L, Donchin E. Performance of concurrent tasks: a psychophysiological analysis of the reciprocity of information-processing resources. Science. 1983;221(4615):1080–2.

32. Cheng KS, Chang YF, Han RP, Lee PF. Enhanced conflict monitoring via a short-duration, video-assisted deep breathing in healthy young adults: an event-related potential approach through the Go/NoGo paradigm. PeerJ. 2017;5:e3857.

33. Schoenberg PL, Hepark S, Kan CC, Barendregt HP, Buitelaar JK, Speckens AE. Effects of mindfulness-based cognitive therapy on neurophysiological correlates of performance monitoring in adult attention-deficit/hyperactivity disorder. Clinical Neurophysiology. 2014;125(7):1407–16.

34. Schoenberg PL, Speckens AE. Modulation of induced frontocentral theta (Fm-θ) event-related (de-) synchronisation dynamics following mindfulness-based cognitive therapy in major depressive disorder. Cognitive neurodynamics. 2014;8(5):373–88.

35. Schoenberg PL, Speckens AE. Multi-dimensional modulations of α and γ cortical dynamics following mindfulness-based cognitive therapy in Major Depressive Disorder. Cognitive neurodynamics. 2015;9(1):13–29.

36. Sanger KL, Dorjee D. Mindfulness training with adolescents enhances metacognition and the inhibition of irrelevant stimuli: Evidence from event-related brain potentials. Trends in Neuroscience and Education. 2016;5(1):1–11.

37. Quaglia JT, Goodman RJ, Brown KW. Trait mindfulness predicts efficient top‐down attention to and discrimination of facial expressions. Journal of personality. 2016;84(3):393– 404.

38. Di Russo F, Martínez A, Hillyard SA. Source analysis of event-related cortical activity during visuo-spatial attention. Cerebral cortex. 2003;13(5):486–99.

39. Fu S, Caggiano DM, Greenwood PM, Parasuraman R. Event-related potentials reveal dissociable mechanisms for orienting and focusing visuospatial attention. Cognitive Brain Research. 2005;23(2-3):341–53.

40. Kelly SP, Gomez-Ramirez M, Foxe JJ. Spatial attention modulates initial afferent activity in human primary visual cortex. Cerebral cortex. 2008;18(11):2629–36.

41. Martinez A, Anllo-Vento L, Sereno MI, Frank LR, Buxton RB, Dubowitz D, et al. Involvement of striate and extrastriate visual cortical areas in spatial attention. Nature neuroscience. 1999;2(4):364.

42. Koenig T, Kottlow M, Stein M, Melie-García L. Ragu: a free tool for the analysis of EEG and MEG event-related scalp field data using global randomization statistics. Computational Intelligence and Neuroscience. 2011;2011:4.

43. Hong X, Wang Y, Sun J, Li C, Tong S. Segregating Top-Down Selective Attention from Response Inhibition in a Spatial Cueing Go/NoGo Task: An ERP and Source Localization Study. Scientific reports. 2017;7(1):9662.

44. Boisgueheneuc Fd, Levy R, Volle E, Seassau M, Duffau H, Kinkingnehun S, et al. Functions of the left superior frontal gyrus in humans: a lesion study. Brain. 2006;129(12):3315–28.

45. Bailey NW, Hoy KE, Maller JJ, Segrave RA, Thomson R, Williams N, et al. An exploratory analysis of go/nogo event-related potentials in major depression and depression following traumatic brain injury. Psychiatry Research: Neuroimaging. 2014;224(3):324–34.

46. Kabat-Zinn J. Wherever you go. There you are: mindfulness meditation in everyday life. 1994.

47. Hergueta T, Baker R, Dunbar GC. The Mini-International Neuropsychiatric Interview (MINI): the development and validation of a structured diagnostic psychiatric interview for DSM-IVand ICD-10. J clin psychiatry. 1998;59(Suppl 20):2233.

48. Steer RA, Beck AT. Beck Anxiety Inventory. 1997.

49. Beck AT, Steer RA, Brown GK. Beck depression inventory-II. San Antonio. 1996;78(2):490–8.

50. Walach H, Buchheld N, Buttenmüller V, Kleinknecht N, Schmidt S. Measuring mindfulness—the Freiburg mindfulness inventory (FMI). Personality and individual differences. 2006;40(8):1543–55.

51. Baer RA, Smith GT, Hopkins J, Krietemeyer J, Toney L. Using self-report assessment methods to explore facets of mindfulness. Assessment. 2006;13(1):27–45.

52. Krompinger JW, Simons RF. Electrophysiological indicators of emotion processing biases in depressed undergraduates. Biological Psychology. 2009;81(3):153–63.

53. Delorme A, Makeig S. EEGLAB: an open source toolbox for analysis of single-trial EEG dynamics including independent component analysis. Journal of neuroscience methods. 2004;134(1):9–21.

54. Palmer JA, Makeig S, Kreutz-Delgado K, Rao BD, editors. Newton method for the ICA mixture model. Acoustics, Speech and Signal Processing, 2008 ICASSP 2008 IEEE International Conference on; 2008: IEEE.

55. Perrin F, Pernier J, Bertrand O, Echallier J. Spherical splines for scalp potential and current density mapping. Electroencephalography and clinical neurophysiology. 1989;72(2):184–7.

56. Tadel F, Baillet S, Mosher JC, Pantazis D, Leahy RM. Brainstorm: a user-friendly application for MEG/EEG analysis. Computational intelligence and neuroscience. 2011;2011:8.

57. Gramfort A, Papadopoulo T, Olivi E, Clerc M. OpenMEEG: opensource software for quasistatic bioelectromagnetics. Biomedical engineering online. 2010;9(1):45.

58. Pascual-Marqui RD. Standardized low-resolution brain electromagnetic tomography (sLORETA): technical details. Methods Find Exp Clin Pharmacol. 2002;24(Suppl D):5–12.

59. Lin F-H, Witzel T, Ahlfors SP, Stufflebeam SM, Belliveau JW, Hämäläinen MS. Assessing and improving the spatial accuracy in MEG source localization by depth-weighted minimum-norm estimates. Neuroimage. 2006;31(1):160–71.

60. Malinowski P, Moore AW, Mead BR, Gruber T. Mindful aging: the effects of regular brief mindfulness practice on electrophysiological markers of cognitive and affective processing in older adults. Mindfulness. 2017;8(1):78–94.

61. Michel CM, Murray MM, Lantz G, Gonzalez S, Spinelli L, de Peralta RG. EEG source imaging. Clinical neurophysiology. 2004;115(10):2195–222.

62. Benjamini Y, Hochberg Y. Controlling the false discovery rate: a practical and powerful approach to multiple testing. Journal of the royal statistical society Series B (Methodological). 1995:289–300.

63. Cramer AO, van Ravenzwaaij D, Matzke D, Steingroever H, Wetzels R, Grasman RP, et al. Hidden multiplicity in exploratory multiway ANOVA: Prevalence and remedies. Psychonomic Bulletin & Review. 2016;23(2):640–7.

64. Lehmann D, Ozaki H, Pal I. EEG alpha map series: brain micro-states by space-oriented adaptive segmentation. Electroencephalography and clinical neurophysiology. 1987;67(3):271–88.

65. Koenig T, Stein M, Grieder M, Kottlow M. A tutorial on data-driven methods for statistically assessing ERP topographies. Brain topography. 2014;27(1):72–83.

66. Murray MM, Brunet D, Michel CM. Topographic ERP analyses: a step-by-step tutorial review. Brain topography. 2008;20(4):249–64.

67. Foxe JJ, Simpson GV. Flow of activation from V1 to frontal cortex in humans. Experimental brain research. 2002;142(1):139–50.

68. Behrmann M, Geng JJ, Shomstein S. Parietal cortex and attention. Current opinion in neurobiology. 2004;14(2):212–7.

69. Malhotra P, Coulthard EJ, Husain M. Role of right posterior parietal cortex in maintaining attention to spatial locations over time. Brain. 2009;132(3):645–60.

70. Hartmann L, Sallard E, Spierer L. Enhancing frontal top-down inhibitory control with Go/NoGo training. Brain Structure and Function. 2016;221(7):3835–42.

71. Wessel JR. Prepotent motor activity and inhibitory control demands in different variants of the go/no‐go paradigm. Psychophysiology. 2018;55(3):e12871.

72. Bruin K, Wijers A. Inhibition, response mode, and stimulus probability: a comparative event-related potential study. Clinical Neurophysiology. 2002;113(7):1172–82.

73. Foxe JJ, Simpson GV, Ahlfors SP. Parieto‐occipital∼ 1 0Hz activity reflects anticipatory state of visual attention mechanisms. Neuroreport. 1998;9(17):3929–33.

74. Fu K-MG, Foxe JJ, Murray MM, Higgins BA, Javitt DC, Schroeder CE. Attention-dependent suppression of distracter visual input can be cross-modally cued as indexed by anticipatory parieto–occipital alpha-band oscillations. Cognitive Brain Research. 2001;12(1):145–52.

75. Hopf J-M, Mangun G. Shifting visual attention in space: an electrophysiological analysis using high spatial resolution mapping. Clinical neurophysiology. 2000;111(7):1241– 57.

76. Worden MS, Foxe JJ, Wang N, Simpson GV. Anticipatory biasing of visuospatial attention indexed by retinotopically specific-band electroencephalography increases over occipital cortex. J Neurosci. 2000;20(RC63):1–6.

77. Foxe JJ, Simpson GV, Ahlfors SP, Saron CD. Biasing the brain’s attentional set: I. Cue driven deployments of intersensory selective attention. Experimental Brain Research. 2005;166(3-4):370–92.

78. Schroeder CE, Mehta AD, Foxe JJ. atDeterminants and mechanisms of attentional modulation of neural processing. Front Biosci. 2001;6:D672–D84.

79. Júnior RdM, Marinho de Sousa B, Fukusima S. Hemispheric specialization in face recognition: From spatial frequencies to holistic/analytic cognitive processing. Psychology & Neuroscience. 2014;7(4):503.

80. Vogel EK, Woodman GF, Luck SJ. Pushing around the locus of selection: Evidence for the flexible-selection hypothesis. Journal of cognitive neuroscience. 2005;17(12):1907– 22.

81. Keil A, Stolarova M, Moratti S, Ray WJ. Adaptation in human visual cortex as a mechanism for rapid discrimination of aversive stimuli. Neuroimage. 2007;36(2):472–9.

82. Eldar S, Yankelevitch R, Lamy D, Bar-Haim Y. Enhanced neural reactivity and selective attention to threat in anxiety. Biological Psychology. 2010;85(2):252–7.

83. Goyal M, Singh S, Sibinga EM, Gould NF, Rowland-Seymour A, Sharma R, et al. Meditation programs for psychological stress and well-being: a systematic review and meta-analysis. JAMA internal medicine. 2014;174(3):357–68.

84. Lutz A, Slagter HA, Rawlings NB, Francis AD, Greischar LL, Davidson RJ. Mental training enhances attentional stability: neural and behavioral evidence. Journal of Neuroscience. 2009;29(42):13418–27.

85. Rauss KS, Pourtois G, Vuilleumier P, Schwartz S. Attentional load modifies early activity in human primary visual cortex. Human brain mapping. 2009;30(5):1723–33.

86. Kerr CE, Jones SR, Wan Q, Pritchett DL, Wasserman RH, Wexler A, et al. Effects of mindfulness meditation training on anticipatory alpha modulation in primary somatosensory cortex. Brain research bulletin. 2011;85(3-4):96–103.

87. Koenig T, Melie-García L. A method to determine the presence of averaged event-related fields using randomization tests. Brain topography. 2010;23(3):233–42.

88. Sibalis A, Milligan K, Pun C, McKeough T, Schmidt LA, Segalowitz SJ. An EEG Investigation of the Attention-Related Impact of Mindfulness Training in Youth With ADHD: Outcomes and Methodological Considerations. Journal of attention disorders. 2017:1087054717719535.

89. Mitchell DJ, McNaughton N, Flanagan D, Kirk IJ. Frontal-midline theta from the perspective of hippocampal “theta”. Progress in neurobiology. 2008;86(3):156–85.

